# Evolution of moss leaf-like organs through variations in deeply conserved developmental principles

**DOI:** 10.64898/2026.02.18.706666

**Authors:** Wenye Lin, Loann Collet, Laure Mancini, Mandar Deshpande, Brendan Lane, Benjamin P. Lapointe, Agnieszka Bagniewska-Zadworna, Anne-Lise Routier-Kierzkowska, Richard S. Smith, Yoan Coudert, Daniel Kierzkowski

## Abstract

Leaves and leaf-like organs with laminar structures and determinate growth arose multiple times independently in land plants. The cellular basis of leaf development is well characterized in flowering plants, and molecular studies have shown that the plant hormone auxin plays a central role in this process, orchestrating cellular growth and differentiation. Auxin is also crucial for the formation of phyllids, the leaf-like organs of bryophytes, yet its precise role in morphogenesis remains unclear. More broadly, whether similar developmental principles are shared across distantly related evolutionary lineages is unknown. Here, we combine live-imaging, genetics, pharmacological treatments, and modeling to investigate the cellular and molecular basis of phyllid development in the model moss *Physcomitrium patens*. By tracking phyllid morphogenesis from a single initial cell to full maturity, we uncover the cellular growth dynamics underlying organ development. We demonstrate that auxin spatially inhibits cell divisions and promotes cellular elongation and differentiation. However, unlike in vascular plants, moss PIN transporters do not participate in polar auxin transport during phyllid development but mainly reduce intracellular auxin concentration. These findings indicate that while auxin’s role in organogenesis is conserved, its transport mechanisms have diverged across land plants. Overall, our study reveals shared principles of planar organ morphogenesis, highlighting how the repeated deployment of similar developmental strategies, with lineage-specific variations, drove the convergent evolution of leaves and leaf-like organs.

**One-Sentence Summary:** Auxin modulates a conserved developmental program in moss leaf-like organs by spatiotemporally controlling cell division and growth.

## INTRODUCTION

Leaves are specialized photosynthetic structures that evolved multiple times independently in the sporophyte generation of vascular plants (*1,2*). They range from large, structurally complex organs, referred to as “true” leaves in flowering plants, to smaller, anatomically simple structures, such as lycophyte microphylls. The repeated emergence of leaves and other leaf-like organs has involved the recruitment of both conserved and distinct genetic mechanisms (*3,4*).

Beyond vascular plants, leaf-like organs known as phyllids evolved convergently in the gametophyte generation of bryophytes, including leafy liverworts and mosses. Phyllids are small lateral appendages that develop at the apex of leafy shoots called gametophores. In the model moss species *Physcomitrium patens*, phyllids have a simple structure, consisting primarily of a single cell layer with a central, multilayered vein – the midrib (*5*). They originate from a single phyllid initial cell that undergoes multiple rounds of oblique divisions, followed by further divisions occurring in its daughter cells (*6*). Despite their immense morphological diversity, and the anatomical differences between bryophyte phyllids and vascular plant leaves, they typically exhibit a laminar structure and determined growth – key adaptations for optimizing light capture and ensuring a similar physiological function.

The molecular basis and cellular dynamics of leaf development have been primarily researched in flowering plants (*7,8*). Studies across diverse species suggest that, despite differences in final morphology, leaf formation follows a shared developmental program that includes common growth dynamics and regulatory mechanisms (*9–12*). For instance, most leaves exhibit gradients of cellular growth and divisions along their proximodistal axis, which are coupled with the progression of cell differentiation (*13–16*). Tuning this common developmental regulation is crucial for the establishment of the final shape and size of the organ (*12,17–19*). While the mechanisms behind these cellular gradients remain unclear, computational modelling and surgical experiments suggest that they may be controlled by unknown intrinsic signal likely diffusing from the base of the organ and creating concentration gradient along its main axis (*12, 14, 16, 17*). These gradients of cell growth and differentiation are modulated by various molecular players such as TCP proteins and miRNAs (*11, 15, 20*), but they may also be at least partially, tuned by plant hormones (*11, 12, 19–21*).

Auxin plays a particularly important role in regulating developmental gradients, as it controls cell division, elongation, and differentiation in a dose-, position- and context-dependent manner (*22*). In angiosperm leaves, local auxin accumulation, driven by polar auxin transport, triggers organ initiation and outgrowth (*23–25*). Auxin maxima at the leaf tip and margin have been proposed to function as global or local organizers, coordinating growth within the developing organ (*12, 17, 26–28*). Subsequently, auxin spreads basipetally from the tip to the base through the leaf epidermis and margin, correlating with the progression of cell differentiation (*19, 28, 29*).

Unlike flowering plants, moss phyllid development is thought to be regulated primarily by a cell lineage-dependent mechanism (*6, 30*). However, cell divisions in the phyllid persist longest at its base (*6, 31, 32*), suggesting that positional cues might also contribute to phyllid organogenesis. Despite the independent evolution origins of moss phyllids and angiosperm leaves, the core genetic components involved in auxin biosynthesis, transport, and signaling are conserved across both lineages, indicating that auxin might provide positional information during phyllid development (*33–38*). However, the spatial and temporal cellular dynamics underpinning phyllid organogenesis remain poorly understood, and it is unknown whether auxin regulates growth and differentiation gradients in this context.

Here, we address these open questions in the model moss *Physcomitrium patens* through a novel time-lapse imaging protocol combined with pharmacological, genetic, and computational approaches. Specifically, we aim to test whether the core morphogenetic principles governing leaf development in flowering plants are shared with bryophytes and contribute to phyllid morphogenesis.

## RESULTS

### Positional cues control growth and cell division during phyllid morphogenesis

The mature upper *Physcomitrium* phyllid is a small (about 2 mm long) laminar organ with a lanceolate shape that is slightly folded along its central midrib (Fig. 1A). Its initiation and early development occur at the apex of the leafy shoot from a single initial cell that is tightly covered by older primordia and numerous axillary hairs. For these reasons, live imaging of the upper phyllid development has been considered too challenging (*6*), which hindered our understanding of the cellular dynamics underpinning its formation and expansion. To overcome these limitations, we developed new dissection and imaging methods allowing us to follow the development of the entire organ from a single initial cell until the phyllid reaches its full size, 5–6 days after initiation (see Methods). Using MorphoGraphX software (*39*), we segmented all cells in 2D or 3D for all time points, and computed cellular growth rates and orientations, tracked cell divisions, and obtained lineage information (Fig. 1, fig. S1, and Movie S1 and S2).

**Fig. 1.**
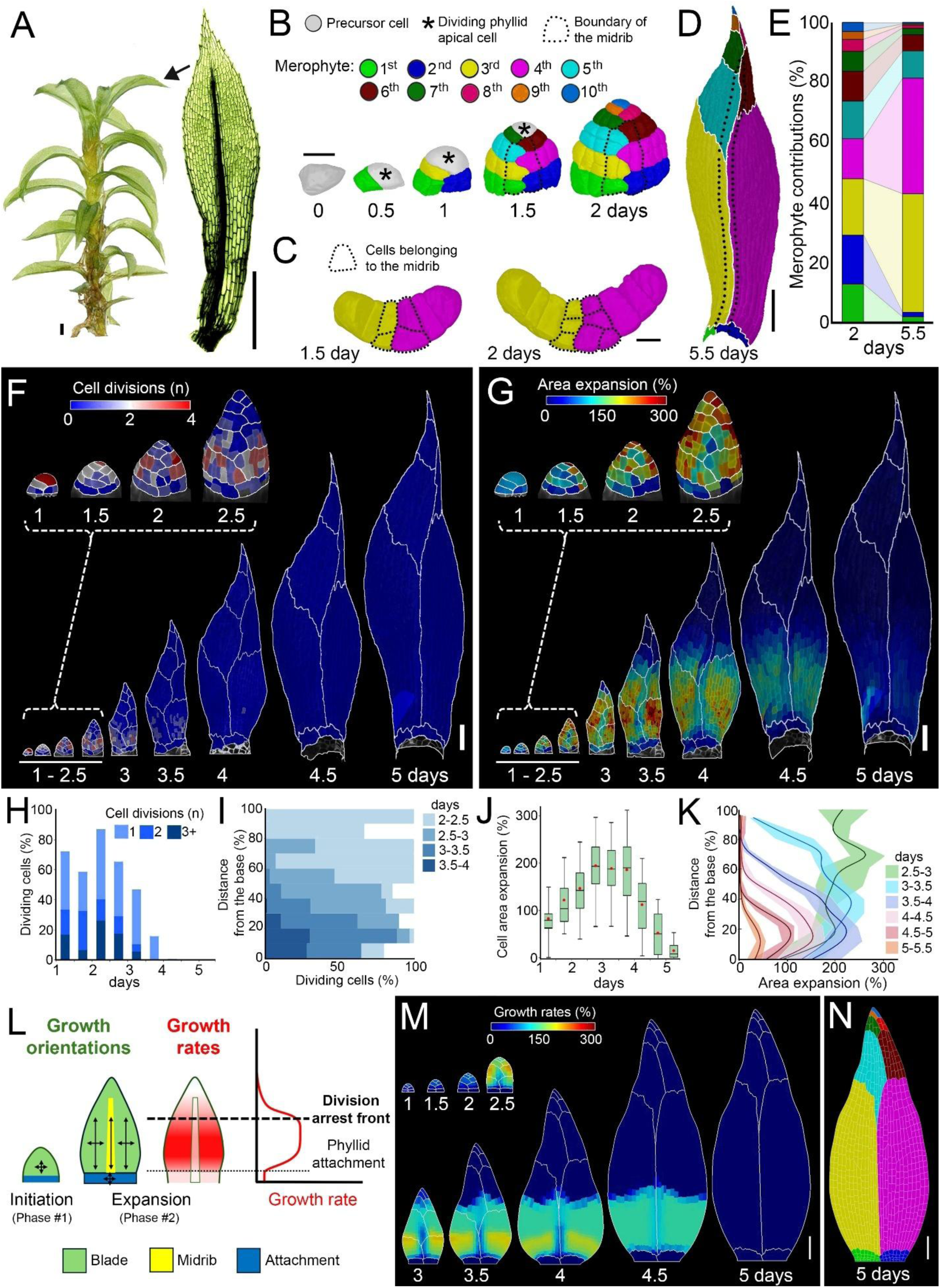
Positional cues control growth and cell division during phyllid morphogenesis. **(A)** *Physcomitrium patens* gametophore isolated from a one-month-old colony and the representative upper phyllid with visible midrib. **(B)** 3D lineage tracing of the upper phyllid initiation. Colors represent merophytes; dotted lines - midrib; asterisks - dividing apical cell. **(C)** Cross-section of the 3^rd^ and 4^th^ merophytes. Dotted lines mark midrib cells. **(D)** Fully developed upper phyllid with colors marking merophytes. **(E)** Relative contribution of different merophytes from 2 days to the final size of phyllid (5.5 days). **(F-G)** Heat maps of cell divisions (F) and area expansion (G) for the upper phyllid. Values are displayed at the earlier time point. **(H-K)** Quantification of cell divisions (H), cell divisions as a function of the normalized distance from the organ base (I), cell area expansion (J), and area expansion as a function of the normalized distance from the organ base (K). The shades contain the second and third quartiles, and the lines indicate the median (n=18, 46, 91, 226, 490, 799, 931, 939, and 938 cells at consecutive time points; three time-lapse series). **(L)** Assumptions of the model of upper phyllid development. Growth is slow and relatively isotropic at initiation (Phase #1). Subsequent growth is homogeneous and anisotropic in the phyllid blade, very anisotropic in the midrib, and isotropic in the attachment zone (Phase #2). Growth rates depend on tissue type and differentiation. Cells stop dividing and differentiate when passing beyond a given distance (thick dotted line) from the phyllid base. Following differentiation (Phase #3), cells stop dividing, elongate, and progressively stop growing. **(M)** Model output colored by areal specified growth rate and merophytes marked with white lines. **(N)** Resultant distribution of merophytes. Scale bars, 500 µm in (A), 20 µm in (C-D), 200 µm in (E), and 100 µm in (G-H and N-O). See also fig. S1 and Movie S1 to S3.

As previously observed, the initiation of the phyllid started with alternating left-right oblique divisions of the precursor cell (Fig. 1B) (*6*, *36*). In the upper phyllid, these divisions consistently generated around 10 daughter cells (10 ± 1 cells, n = 23 primordia) that gave rise to a series of clonal sectors (merophytes) with a zigzagging boundary in the middle of the organ (Fig. 1B, D and E). This zigzagging pattern subsequently straightens as strong elongation of the midrib aligns initially oblique sector boundaries with the main (longitudinal) organ axis (Fig. 1F). The first rounds of subsequent divisions of the daughter cells (from 1-2 days) were always longitudinal (oriented parallel to the main axis of the organ) and produced additional cell files in the basal merophytes (Fig. 1B). Subsequently, a single cell in each sector, located in the middle of the primordium, divided longitudinally, giving rise to the midrib (Fig. 1B and C). These predictable patterns of precursor cell divisions, as well as a consistent number of merophytes, support the idea that phyllid initiation is controlled by lineage-based cues (*6*).

We next evaluated whether merophytes could act as autonomous domains, inheriting developmental instructions from their initials as previously suggested (*6*). To investigate this, we quantified cell growth and divisions throughout phyllid development (Fig. 1F to K, and Movie S1). During the initiation phase (0–2 days), cellular growth was relatively slow and uniform, while most cells continued dividing (Fig. 1, F, H, and J). From day 2, cells in the primordium—except those at the phyllid base, which form attachments to the stem—accelerated their growth along the longitudinal axis of the organ (Fig. 1, G, J, and K, fig. S1, D to H, and Movie S1 and S2). Shortly after midrib establishment (at 2 days), the mediolateral growth in this region became strongly restricted (Fig. 1G, and fig. S1, D and E). From 2.5 days after initiation, cell divisions progressively decreased and a gradient emerged, with divisions gradually ceasing from the tip to the base (Fig. 1, F, H, and I). This was followed by a basipetal gradient of cellular growth, first visible at 3–3.5 days and persisting until the end of phyllid development (Fig. 1, G and K). Prolonged growth at the phyllid base resulted in a substantial size increase in the proximal sectors, with the 3^rd^ and 4^th^ sectors contributing nearly 80% for the final organ size (Fig. 1, D and E). Contrary to expectations of inherited behavior from the phyllid initial cell, cell divisions and growth did not follow patterns that would be expected solely from clonal regulation within merophytes but formed a global gradient irrespective of sector boundaries (Fig. 1, F and G). This suggests that positional cues rather than sector-dependent lineage control phyllid development after initiation. Additionally, both cell divisions and growth were maintained at a relatively fixed distance from the organ base (Fig. 1, F-G, and fig. S1, D-E), indicating a global regulation of organ morphology, possibly mediated by signals originating from the organ base. Taken together, our live-imaging data suggest that phyllid initiation is mainly driven by lineage-autonomous behavior, whereas subsequent development across sectors is primarily regulated by positional cues that spatially restrict cell divisions to the proximal organ region.

To test this hypothesis and determine whether the identified developmental rules were sufficient to account for phyllid shape, we integrated our quantitative observations into a simple 2D computational model of phyllid development. In our model, cells are represented as polygons, and can individually control their growth and cell division depending on their location within the phyllid. The simulation followed three distinct phases of phyllid development (Fig. 1L). Phase #1 or phyllid initiation (0 to 1.5 days): the primordium is divided into an upper zone and a phyllid attachment zone that is composed of undifferentiated cells growing slowly and isotropically. Phase #2 or phyllid expansion (2 to 4 days): cell division and growth accelerate in the upper zone of the phyllid, while the phyllid attachment zone grows very slowly. The midrib is introduced with reduced growth centrally along the mediolateral organ axis. Phase #3 or phyllid maturation (after 4 days): all cell divisions cease, and the phyllid progressively stops expanding. The main assumptions of the model are as follows (see Table S1 for model parameters): (1) Cells have positional information and know their location between the base of the phyllid and the apical cell.

This creates a polarity field. (2) The apical cell divides obliquely in an alternating left-right pattern, and daughter cell divisions follow the shortest of the paths parallel or perpendicular to the polarity field. (3) Undifferentiated, dividing cells grow relatively fast. (4) Cells divide upon reaching a predetermined size threshold. (5) Beyond a given distance from the organ base, cells stop dividing and differentiate. (6) Differentiating cells stop dividing, elongate transiently before gradually ceasing growth as observed in time-lapse data (Fig. S1, A-C). In the model, cells decide their own growth rates depending on their location in the phyllid, and the growth rate is specified at the cellular level.

Although the 2D model includes only a minimal set of roles, we found that it could qualitatively recapitulate the growth dynamics of developing phyllids by using positional cues to control growth and cell division. This resulted in the patterning of sectors, including their relative size and distribution, comparable to those observed experimentally (Fig. 1, D, F, G, M, and N, and Movie S3). This indicates that the minimal set of assumptions (Table S1) is sufficient to explain how phyllids achieve their final shape. Furthermore, this result supports the hypothesis that positional information controls phyllid development.

### Auxin acts as a positional cue, with PINs reducing intracellular auxin concentration to shape phyllid development

Global growth and cell division gradients appear to govern phyllid development, but the nature of underlying signals through which they are controlled remain unclear. During vascular plant organogenesis, auxin plays a critical role in providing positional information for the establishment of tissue patterns, organ polarity, and the coordination of cell differentiation and growth (*12*, *19, 21–28*, *40*). Moreover, alterations in auxin levels or transport affect phyllid shape and size in *Physcomitrium* (*34–37*, *41*), suggesting that this phytohormone may be crucial for regulating gradients of cell division and growth during phyllid morphogenesis.

To investigate whether the role of auxin as a positional cue is conserved in moss organogenesis, we first monitored the dynamics of auxin biosynthesis gene expression and auxin sensing during upper phyllid development, both spatially and temporally. In *P. patens*, auxin is mainly synthesized through the tryptophan-dependent pathway controlled by *TRYPTOPHAN AMINO-TRANSFERASE* (*TAR)* and *YUCCA* (*YUC)* genes (*42, 43*). Published datasets show that *TARA*, *TARC*, and *YUCF* genes are the most highly expressed in phyllids (Fig. S2), consistent with previous reports of similar promoter activity for *TARA* and *TARC* (*44*). We detected a clear GFP signal in phyllids of the *TARA* promoter line, but not in the *YUCF* promoter line. The most parsimonious explanation is that *YUC* expression remains below our reporter’s detection threshold. Because the conversion of TRP to IPyA by TAR enzymes is a limiting step in auxin biosynthesis in *Physcomitrium* (*42*), we reasoned that a *TAR–GFP* reporter would provide a suitable proxy for identifying putative auxin biosynthesis sites in phyllids. The promoter activity of *TARA2* was low on day 1, peaked on day 2, and rapidly declined to undetectable levels by day 3 after upper phyllid initiation (Fig. 2A). TARA promoter activity was first observed in the apical cell and then mainly detected in a few cells at the primordium tip and distal margin (Fig. 2A). These data suggest that auxin is likely produced at the organ tip during the early developmental stages of the upper phyllid. Consistently, auxin sensing monitored by the ratiometric biosensor *R2D2* was first detected in the distal part of the upper phyllid and subsequently spread toward its base (Fig. 2, D and F, fig. S3C), in alignment with a previous report (*45*). Together, these data indicate that auxin moves throughout the organ from its source cells at the phyllid tip toward its base.

**Fig. 2.**
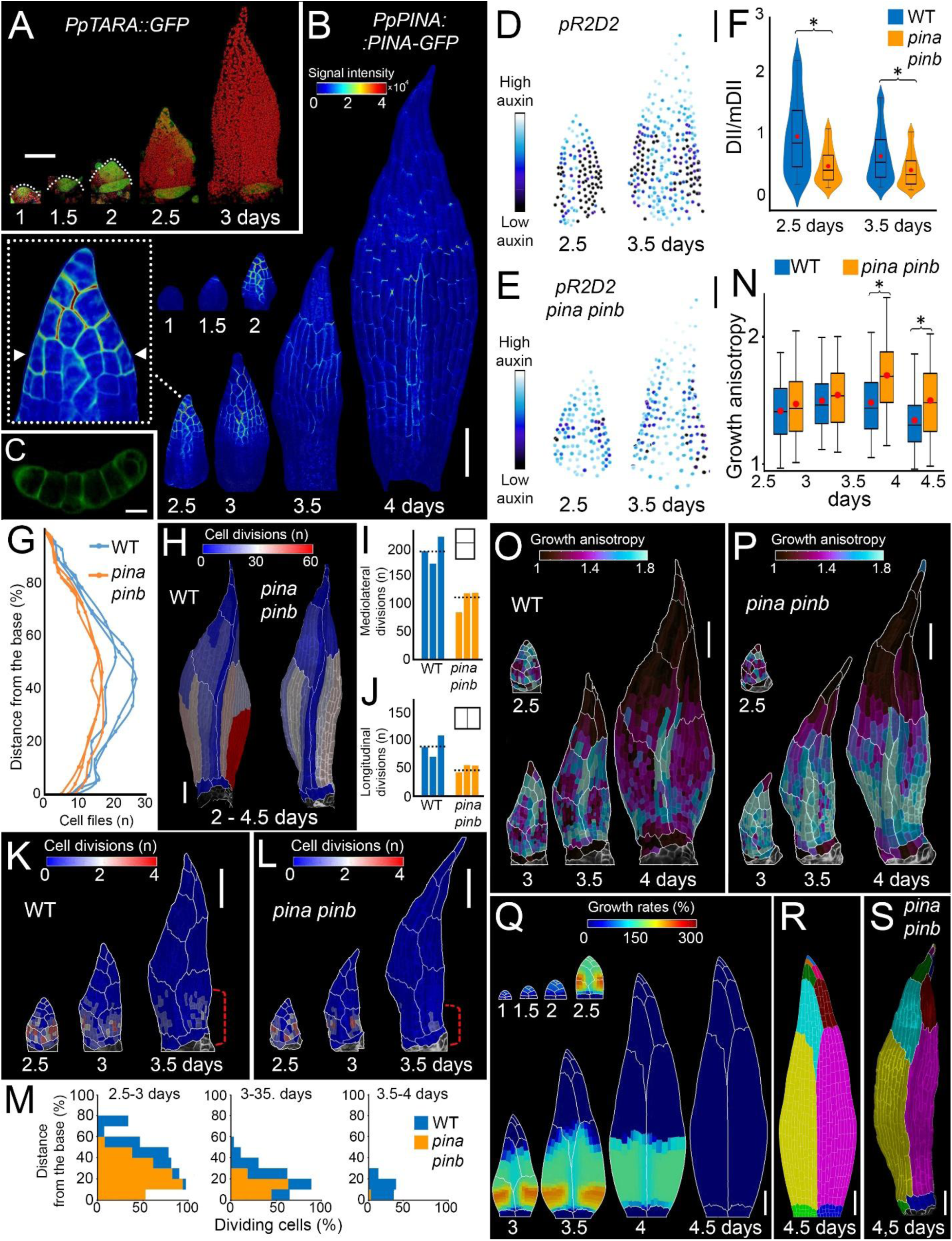
Auxin acts as a positional cue, with PINs reducing intracellular auxin concentration to shape phyllid development. **(A)** Localization of *PpTARA::GFP* (green) expression in the upper phyllid. Chloroplast fluorescence in red. **(B-C)** Localization of *PpPINA::PINA-GFP* in the upper phyllid. Heatmap represents the intensity of PINA-GFP signal. Inset: close-up view of the phyllid. Arrowheads indicate the position of the cross section shown in (C). **(D-E)** Images of segmented nuclei in phyllids at 2.5 and 3.5 days. Each nucleus is color-coded according to the relative level of auxin sensing (DII/mDII signal ratio). **(F)** Quantification of DII/mDII signal ratio, n=3 phyllids for WT and *pina pinb* at 2.5 days, n=5 phyllids for WT and *pina pinb* at 3.5 days. Asterisks indicate statistical significance, P<0.05, t-test. **(G)** Quantification of phyllid width in cell files as a function of the normalized distance from its base. **(H)** Heatmaps of cumulated cell divisions (2-4.5 days) for WT and *pina pinb* upper phyllids. **(I-J)** Quantification of mediolateral (I) and longitudinal (J) cell divisions (2-4.5 days) in WT and *pina pinb*. Dotted lines show average values for three time-lapse series. **(K-L)** Heatmaps of cell divisions in WT (K) and *pina pinb* mutant phyllids (L). Red brackets indicate the extent of the cell division zone. **(M)** Quantification of the number of dividing cells as a function of the normalized distance from the phyllid base. **n.** Quantification of growth anisotropy in WT and *pina pinb* mutant, n=3 time-lapse series for WT and *pina pinb*. Asterisks indicate statistical significance, P<0.05, t-test. **(O-P)** Heatmaps of growth anisotropy in the WT (O) and *pina pinb* mutant (P). Heat values are displayed at the earlier time point. **(Q-R)** Model of the *pina pinb* mutant phyllid with reduced cell divisions and increased growth anisotropy compared to WT. Model output colored by areal specified growth rate (Q). Resultant distribution of merophytes (R). **(S).** Upper phyllid of *pina pinb* mutant with colors marking merophytes. Scale bars, 100 μm (B, D-E, H, K-L, O-P, and Q-S), 10 μm (C), 20 μm (A). See also fig. S2 to S7 and Movies S4 to S7.

In angiosperms, the spatial distribution of auxin within tissues is primarily governed by auxin efflux carriers belonging to the *PIN-FORMED* (*PIN*) family, which are responsible for polar auxin transport (*23–25*, *46, 47*). The role of PIN proteins in exporting auxin out of cells is conserved in *Physcomitrium* (*35*, *48*), whose genome encodes three canonical PINs: *PINA*, *PINB*, and *PINC*. All three genes are expressed in phyllids with both *PINA* and *PINB* genes detected at similar levels, significantly higher than *PINC* (*34*, *38, 49, 50*). While phyllid development is only mildly affected in single mutants, it is clearly disrupted in *pina pinb* double mutant that develops narrower organs, indicating that PINA and PINB act redundantly to control phyllid morphogenesis (*32*, *38*). To understand the role of efflux carriers in the dynamics of auxin transport, we precisely monitored the distribution of PINA and PINB expression during upper phyllid development. PINA-GFP was first detected in the apical cell 1.5 days after organ initiation, and its expression domain subsequently expanded toward the more proximal regions of the phyllid, invading up to two thirds of its length (Fig. 2B). PINB-GFP expression patterns and timing were overall similar to PINA-GFP (Fig. S3 A, B). The basipetal spread of the expression of these transporters and accompanying auxin sensing (*R2D2*) correlated with the progression of the cell division arrest front (Fig. 1G, and Fig. 2K). This suggests that PINs may mediate a basipetal auxin movement to regulate phyllid development through auxin-mediated inhibition of cell divisions.

If this hypothesis is correct, eliminating PIN activity should limit the progression of cell division arrest, and consequently expand the division zone toward the phyllid tip as auxin would no longer travel as far as in the wild-type from its source at the phyllid tip to inhibit divisions. To test this hypothesis, we introduced a plasma membrane marker in a *pina pinb* double mutant (*34*) and monitored the development of the upper phyllid with time-lapse imaging. The *pina pinb* mutant phyllids were composed of fewer cell files than WT phyllids, in agreement with previous studies (Fig. 2G) (*32*). Cell divisions were reduced in both longitudinal (−51% on average) and mediolateral (−43% on average) orientations (Fig. 2, H to J). However, contrary to our expectations, while the general shape of basipetal gradients of cell division were comparable to the wild type, the division zone was spatially reduced to more proximal regions of the phyllid in the mutant (Fig. 2, K to M, fig. S4, and Movie S4). The duration of cell divisions was also shortened in the absence of auxin efflux carriers (Fig. 2M, fig. S4, A and C, and Movie S4). These observations raised the hypothesis that the lack of *PINA* and *PINB* could promote auxin diffusion through the organ by increasing its intracellular concentration, instead of restricting it. To test this, we knocked-out *PINA* and *PINB* in the transgenic line expressing the *R2D2* biosensor using CRISPR/Cas9-mediated gene editing and recovered mutant lines displaying phyllid shape defects similar to those previously reported (fig. S5). Consistently, we observed a significantly lower DII/mDII signal ratio in the *pina pinb* mutant compared to WT, reflecting an overall increase of auxin sensing levels throughout phyllids (Fig. 2, D to F, fig. S3 C).

To explain these observations, we closely examined the subcellular localization pattern of PINA and PINB in growing phyllids. During the early stages of phyllid development, both PINA-GFP and PINB-GFP signals were primarily detected at membranes, without any clear basal polarization (inset in Fig. 2B; fig. S3 A and B). The previously reported polar or bipolar localization at the apical and basal membranes (*35*) was only observed in differentiated, non-growing cells at the tips of the phyllids, starting from 3.5 days after initiation (Fig. 2B). Intriguingly, we found that a portion of PINA and PINB proteins was localized at membranes facing the extracellular space, suggesting that PINs might export auxin from the cytoplasm to the cell wall and potentially out of the organ (Fig. 2C; fig. S3 B). Together, these findings indicate that, in contrast to vascular plants, plasma membrane localized PINs in moss may not participate in basipetal auxin movement during early phyllid development but rather reduce an overall intracellular auxin levels. In their absence, auxin could instead diffuse between cells via plasmodesmata, a scenario consistent with the presence of plasmodesmata in anticlinal walls observed during early phyllid development (fig. S6) (*29*, *51–56*).

Next, to test whether the inhibition of cell division affects organ shape as observed in *pina pinb* phyllid (Fig. 2, G and H), we modified the parameterization of our computational model of the upper phyllid (Fig. 1, M and N) by reducing the distance from the base at which cells are allowed to proliferate, and accelerating the transition from Phase #2 to Phase #3, when all cell divisions cease. We did not aim to explicitly model auxin transport, but rather to evaluate whether auxin-induced changes in cell behavior could account for observed changes in phyllid morphology. These modifications reflect the observed higher auxin spread from the organ tip in the *pina pinb* mutant and assume that elevated auxin levels can override the influence of the basal zone, thereby forcing cells to differentiate earlier. With these simple changes, the model produced a narrower phyllid, resembling that of the *pina pinb* mutant. Nevertheless, the simulated organ was shorter than experimentally observed, suggesting that auxin-mediated inhibition of cell division alone is not sufficient to fully account for the mutant phenotype (fig. S7, and Movie S5).

Apart from accelerating the transition from cell division to differentiation, auxin is also known to control cell elongation (*12*, *41*). Indeed, we observed a significant increase in cellular growth anisotropy in the *pina pinb* double mutant (Fig. 2, N to P, fig. S8, and Movie S6). We tested the impact of this additional parameter by increasing the cell elongation rate in the model and found that the resulting simulated phyllid shape and sector distribution matched those observed in the *pina pinb* mutant (Fig. 2, Q to S, and Movie S7). Altogether, our experiments and simulations show that auxin can fine-tune phyllid shape by modulating the underlying gradients through the inhibition of cell division and promotion of cell elongation.

### Auxin can disrupt cell division and growth gradients during phyllid development

Auxin appears to be crucial for phyllid morphogenesis by promoting the cessation of cell divisions and stimulating cell elongation, thereby influencing the organ’s final shape. When the auxin concentration is increased—such as through the removal of PIN proteins—higher auxin reduces the proliferative activity (Fig. 2, K-M) and increases growth anisotropy (Fig. 2, N-P). This suggests that auxin is a major regulatory cue, sufficient to counteract intrinsic basal gradients that would otherwise sustain cell division at the phyllid base. To investigate this further, we sought to test whether auxin had the capacity to globally alter phyllid development by reprogramming cell behaviour across the entire organ. We therefore hypothesized that supplying exogenous auxin should lead to a rapid cessation of cell divisions and enhanced cell elongation throughout the primordium, ultimately producing a narrow phyllid composed of fewer, elongated cells.

To explore this idea, we first used our model to simulate the growth of the upper phyllid under overall elevated auxin levels. Specifically, the transition from Phase#2 to Phase#3 was accelerated (*i.e.*, faster cessation of cell divisions) and cell elongation was increased, compared with the wild-type phyllid simulations (Table S1). Under these assumptions, our model produced a very narrow, elongated phyllid (Fig. 3A, and Movie S8).

**Fig. 3.**
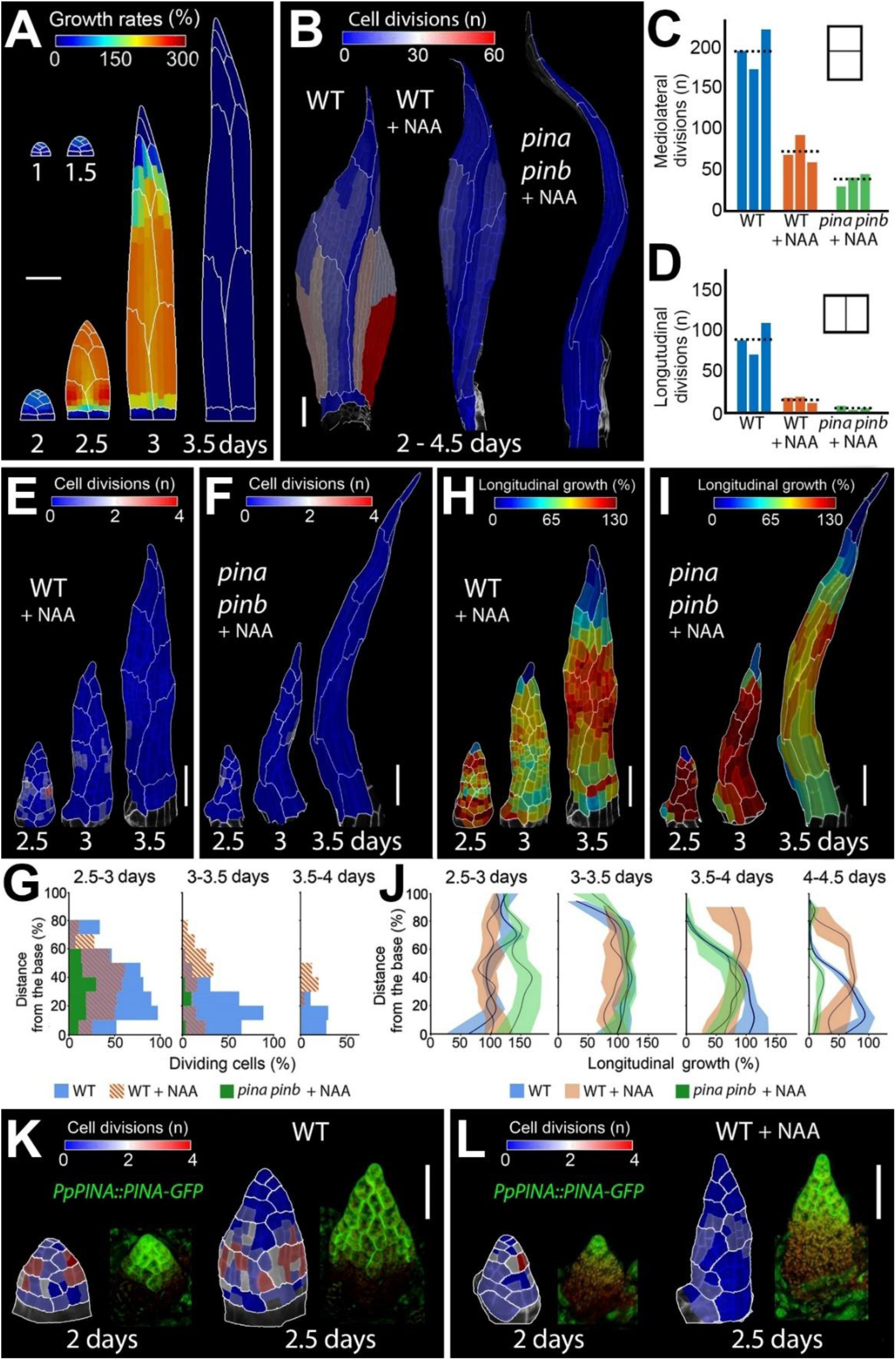
Auxin can disrupt the basipetal gradients during phyllid development. **(A)** Model of the phyllid with cell divisions eliminated after Phase #1 and an increased growth anisotropy. Model output colored by areal specified growth rate. **(B)** Heatmaps of cumulated cell divisions (2-4.5 days) for WT and *pina pinb* mutants upper phyllids treated with auxin. **(C-D)** Quantification of mediolateral (C) and longitudinal (D) cell divisions (2-4.5 days) in WT and *pina pinb* phyllids treated with auxin (three time lapse series). Dotted lines show average values. **(E-F)** Heatmaps of cell divisions in WT (E) and *pina pinb* mutant (F) treated with auxin. **(G)** Quantification of the number of dividing cells as a function of the normalized distance from the phyllid base. **(H-I)** Heatmaps of longitudinal growth in WT (H) and *pina pinb* mutant phyllids (I) treated with auxin. **(J)** Quantification of the longitudinal growth as a function of the normalized distance from the phyllid base (226, 490, 799, 931 cells for WT, 193, 271, 321, 325 cells for WT treated with auxin, and 201, 234, 239, 237 cells for *pina pinb* treated with auxin; three time-lapse series). Heat values are displayed at the earlier time point. **(K-L)** Localization of the cell divisions and the expression of *pPINA::PINA-GFP* at early developmental stages of upper phyllids in the WT (K) and WT treated with auxin (L). Heat maps of cell divisions (left) and PINA-GFP localization (right). GFP in green, autofluorescence in red. Scale bars, 100 μm (A-B, E-F, and H-I) and 50 μm (K-L). See also fig. S8 to S12 and Movies S8 to S12.

To validate the model’s predictions, we treated one-day-old upper phyllid primordia with 1-naphthaleneacetic acid (NAA), a PIN-transportable synthetic auxin, and tracked their subsequent development with time-lapse imaging. NAA treatment significantly altered phyllid morphology. As expected, cell divisions and mediolateral growth were reduced, while longitudinal growth rates increased, producing much narrower organs compared to both the wild-type and the *pina pinb* mutant (Fig. 3, B to D, fig. S9, A, C, E, and fig. S10). Unexpectedly, however, cell divisions did not cease uniformly across the organ. Instead, active cell division persisted in the central region of the phyllid, whereas the basal region—which typically exhibits high division rates—stopped dividing earlier (Fig. 3, E and G, fig. S9, A, C, and E, and Movie S9). The typical growth gradient was also disrupted, with cell growth lasting longer in the middle than at the base (Fig. 3, H and J, fig. S6, B, D, and F, fig. S8, and Movies S9 and S10). The sustained mediolateral growth in the organ’s center (fig. S8, and Movie S10) ultimately led to the formation of a lanceolate phyllid, but not a very narrow organ as predicted by our model (Fig. 3, A and B).

This raised the question of why cell divisions and growth continued in the middle region of NAA-treated phyllids. Our data suggested that intracellular auxin levels decrease due to the activity of PIN proteins (Fig. 2, D to F). We therefore hypothesized that regional differences in PIN expression could determine how cells respond to exogenous auxin. In wild-type plants, PINA expression was restricted to the organ tip—where cells elongate and differentiate—but also partially overlapped with the distal region of the division zone (Fig. 2B, and Fig. 3K). Instead, PINA expression was absent at the phyllid base, where most cell divisions take place (Fig. 2B, and Fig. 3K). Upon NAA treatment, we observed that this apical-basal distribution pattern remained largely unchanged (Fig. 3L, and fig. S11). From these observations, we speculated that when exogenous auxin is applied, cells located at the phyllid base are likely to experience higher auxin levels and thus cease dividing first. Conversely, in the upper portion of the proliferative zone (adjacent to or overlapping with PINA expression), PIN-mediated auxin efflux may prevent cells from reaching the auxin threshold required for immediate differentiation.

If this is correct, then external auxin application in the absence of PINs should further eliminate the divisions in this middle zone. Consistent with this hypothesis, treating the *pina pinb* mutant with NAA caused cell divisions to cease very early and largely abolished the sustained divisions observed in the central region of auxin-treated wild-type phyllids (Fig. 3, B to G, fig. S12, and Movie S11). In addition, longitudinal growth was further enhanced while mediolateral growth reduced, effectively canceling the inverted growth gradient observed in the NAA-treated wild-type (Fig. 3, H to J, fig. S13, and Movie S12). As a result, a very narrow, rod-like phyllid developed, resembling the model outcome predicted for auxin-treated phyllids (Fig. 3A).

Altogether, these data show that auxin is sufficient to disrupt intrinsic global gradients of cell division and growth. This observation further supports the idea that PINs in moss can locally lower auxin concentration to regulate the switch from cell divisions to elongation. This result highlights the importance of PIN-driven auxin homeostasis in shaping phyllid development, underscoring a complex interplay between global auxin gradients and localized PIN activity in determining the spatial patterns of cell division and elongation.

### A temporal shift in auxin-triggered cell differentiation accounts for the juvenile-to-adult phyllid shape transition

Phyllid morphology changes gradually during *Physcomitrium* gametophore ontogeny (Fig. 4A) (*31*, *32*), a process known as heteroblastic development (*57*). Basal (juvenile) phyllids lack a midrib and are substantially smaller, containing significantly fewer cells than upper phyllids (Fig. 4, A to C). They consistently maintain a narrow width, except at the tapering tip (Fig. 4A). This narrow morphology with a reduced cell number, mirrors that of auxin-treated upper (adult) phyllids, especially in the *pina pinb* mutant (Fig. 3B), suggesting that auxin could play a role in the juvenile-to-adult phase transition in moss.

**Fig. 4.**
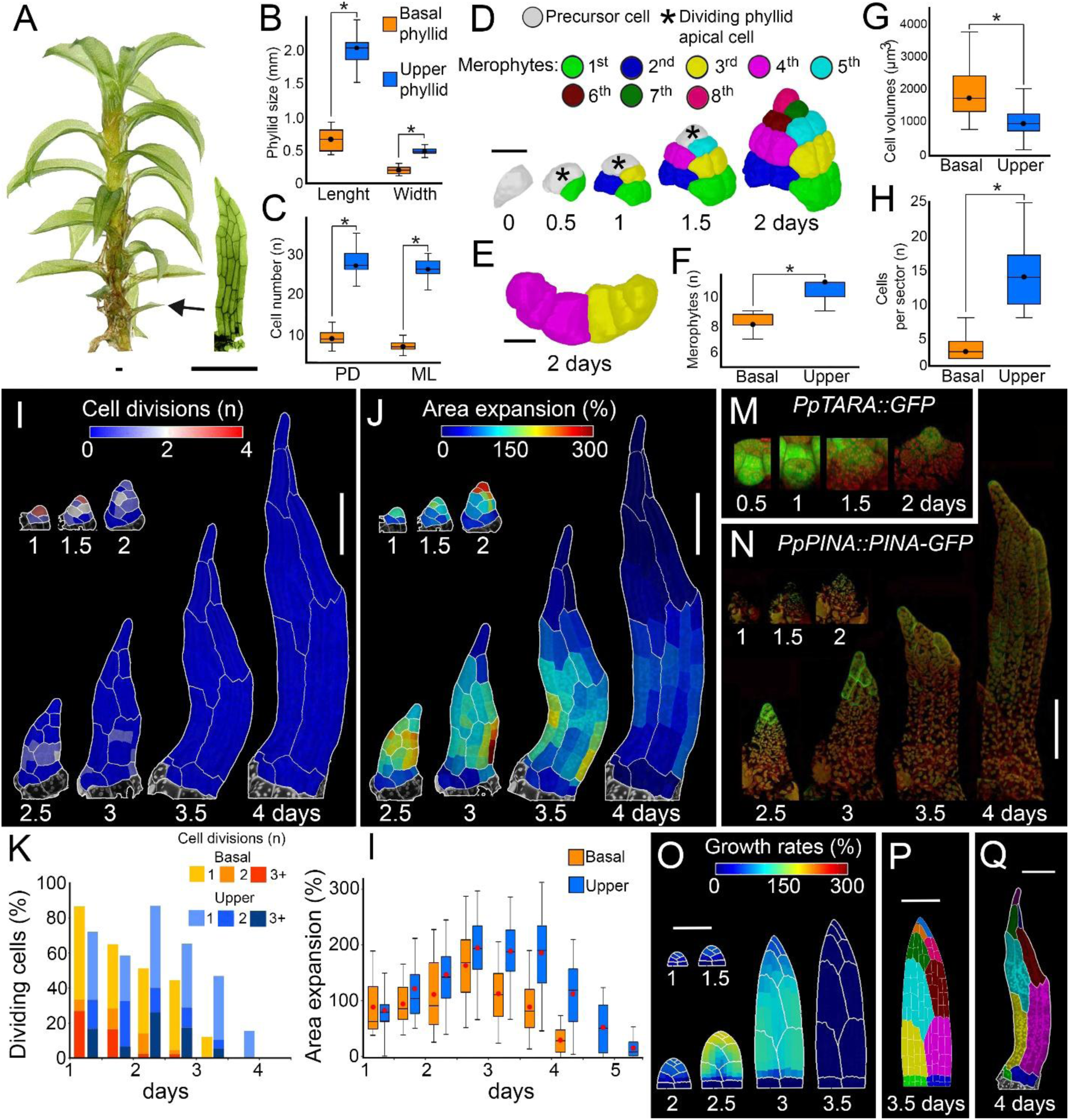
A temporal shift in auxin-triggered cell differentiation accounts for the juvenile-to-adult phyllid shape transition. **(A)** *Physcomitrium patens* gametophore isolated from a one-month-old colony and a representative basal phyllid. **(B-C)** Quantification of basal (juvenile) and upper (adult) phyllid size (B) and cell number (C) along proximodistal (PD) and mediolateral (ML) organ axes (n=21 and 35 basal and upper phyllids respectively). **(D)** 3D lineage tracing of the basal phyllid initiation. Colors indicate merophytes and asterisks dividing apical cells. **(E)** Cross-section of the 3rd and 4th merophytes. **(F-H)** Quantification of the number of merophytes per organ (F) (n=26 phyllids in both basal and upper), cell volumes in 3rd, 4th, and 5th merophytes (n=4 basal and upper phyllids) (G), and number of cells per major merophyte at 2 days (n=12 and 12 basal and upper phyllids) (H). **(I-J)** Heat maps of cell divisions (I) and area expansion (J) for the basal phyllid. Values are displayed at the earlier time point. **(K-L)** Quantification of cell divisions (K), and cell area expansion (L) (n=15, 43, 94, 158, 238, 201 and 205 cells at consecutive time points for the basal phyllid and 18, 46, 91, 226, 490, 799, 939 and 938 for the upper phyllid; three time-lapse series). **(M-N)** Localization of *PpTARA::GFP* (M) and *PpPINA::PINA-GFP* (N) in the basal phyllid. GFP in green, chloroplast fluorescence in red. **(O-P)** Model of the basal phyllid where cell division stops earlier as compared to the upper phyllid. Model output colored by areal specified growth rate (O). Resultant distribution of merophytes (P). **(Q)** Basal phyllid with colors marking merophytes. Asterisk indicates significance with P < 0.05, t-test. Scale bars, 500 µm in (A), 20 µm in (D and F), and 100 µm in (I-J, and O-Q). See also figs. S13, 14 and S15, and Movies S13 to S15.

To investigate this, we first analyzed when the initial differences between basal and upper phyllids were established. Although the alternating left-to-right oblique divisions of the precursor cell were observed in both basal and upper organs (Fig. 1B, and Fig. 4D), the apical cell produced significantly fewer merophytes in the basal phyllid (8±1 vs. 10 ± 1 cells) (Fig. 4, D to F).

Merophytes also consisted of fewer but bigger cells as early as 2 days after initiation, and longitudinal divisions were completely absent in the basal phyllid (Fig. 1C, and Fig. 4, F to H). Additionally, cell divisions and cell growth stopped around 1 day earlier in the basal phyllid (Fig. 4, I to L, and Movie S13) as compared to the upper phyllid (Fig. 1, F to G, and Movie S1). This timing was similar to what we observed upon auxin treatment of the upper phyllid in the *pina pinb* mutant (Fig. 3, F and G). Although basipetal gradients of cell division and expansion were present in the basal phyllid, they persisted very briefly (Fig. 4, I and J, and Movie S13). After initiation, the longitudinal growth was comparable to that observed in the upper phyllid, while the mediolateral growth was strongly restricted (fig. S14, and Movie S14). Altogether this indicates that a very early onset of cell differentiation in the basal phyllid, rather than changes in the basipetal cell division gradient shape, controls heteroblastic development in the moss.

To test the role of auxin in driving this early shift from cell division to differentiation, we first examined auxin biosynthesis in the basal phyllid using a *TARA* promoter reporter line (*43, 44*). Unlike in the upper phyllid, where *TARA* expression peaked at 2 days post-initiation (Fig. 2A), the basal phyllid showed a strong *TARA* signal in the initial cell at 0.5 days, which gradually diminished until 2 days after initiation (Fig. 4M). This finding suggests that auxin levels might be higher during the very early stages of basal phyllid outgrowth, compared with the upper phyllid. Additionally, both PINA-GFP and PINB-GFP expressions in the basal phyllid were much weaker than in the upper phyllid, appearing only in a few differentiating cells at the organ tip from 2.5 days post-initiation before fading rapidly (Fig. 4N and fig. S15), which indicated a reduced auxin export capacity. Unfortunately, technical limitations prevented us from obtaining reliable R2D2 signal measurements in developing basal phyllids. Collectively, these observations support the idea that an early, auxin-triggered transition from cell divisions to differentiation may underlie basal phyllid morphology.

We tested this further using model simulations. Specifically, we modified our initial model of upper phyllid development (Fig. 1, M and N) by accelerating the timing of cell division cessation (*i.e.*, a faster transition from Phase #2 to Phase #3). With this simple change, the model simulated a smaller organ, similar in shape and size to the observed basal phyllid (Fig. 4O, and Movie S15). In contrast to the model of the upper phyllid, this simulation produced a narrow organ with merophytes located near its base contributing much less to overall organ size, which reflected our biological observations (compare Fig. 1, D and N with Fig. 4, P and Q). Altogether, these data indicate that a temporal shift in the auxin-driven basipetal cell differentiation gradient is likely sufficient to account for the shape changes characterizing the juvenile-to-adult phase transition, a defining feature of phyllid heteroblastic development.

## DISCUSSION

This study sought to elucidate the fundamental principles underlying phyllid morphogenesis in the moss *Physcomitrium patens*, a model bryophyte. By applying innovative methods to track and quantify cellular growth from a single initial cell to a fully expanded organ, we characterized phyllid development at an unprecedented spatiotemporal resolution.

Our data confirms previous observations that lineage-based cues dictate the division patterns of apical cell derivatives during the onset of phyllid development (*6*). However, subsequent stages appear to be mainly regulated by global gradients of cell division, growth, and differentiation, independent of cell lineage. Cell divisions are maintained at a defined distance from the base of the organ and gradually cease following a basipetal wave of differentiation. Both upper and basal phyllids follow a similar developmental trajectory, but the smaller size of basal phyllids results from an earlier cessation of cell division and a premature transition to cell differentiation.

Overall, phyllid morphogenesis parallels key aspects of leaf development previously described in *Arabidopsis thaliana*. While early *Arabidopsis* leaf patterning is guided by positional information—in contrast to moss phyllids—later developmental stages feature a basipetal gradient of cell division and growth, with marked differences in the timing of gradient establishment in juvenile and adult organs in both plant lineages (*6*, *12*, *16*, *18*, *19*). Moreover, cell division and growth zones are restricted to the organ base, suggesting a conserved role for basal intrinsic cues in patterning these zones. Consistent with our findings in moss phyllids, a model of leaf morphogenesis supports a key role for basally derived signals, though their molecular nature remains unclear (*12*, *16*, *17*). Beyond *Arabidopsis*, comparative morphometric analyses across eudicots indicate that similar leaf shapes can be attained through diverse late growth patterns. While the early phase of cell divisions is generally conserved, the directionality of differentiation may vary, with some species exhibiting acropetal or bidirectional waves, or a more even distribution of both divisions and differentiation (*15*).

In angiosperm organs, developmental gradients are closely linked to the dynamics of the phytohormone auxin. For instance, auxin promotes cell differentiation and growth in leaves (*11*, *12*), and the spatial distribution of auxin responses and transporters correlate with the progression of cell differentiation in both leaves and floral organs (*19*, *29, 58*, *59*). Through genetic perturbations and pharmacological assays, we provide multiple lines of evidence that auxin plays similar roles in *Physcomitrium* phyllid development. Specifically, auxin likely synthesized at the phyllid tip serves as a spatiotemporal regulator of global gradients of cell division and growth, and it specifically promotes the transition from cell division to cell elongation and differentiation (*44*). By locally reducing intracellular auxin levels, evolutionarily conserved plasma membrane-localized PIN proteins fine-tune phyllid shape and size (*34*, *35*). Under normal conditions, auxin produced at the upper phyllid tip moves basipetally to induce cell differentiation, while cell divisions remain restricted to the organ base. PINA and PINB are frequently present at the membranes facing the outside of the organ in both phyllid tip, margin, and midrib (Fig. 2C; figs S 3A and B; fig. S15). Such localization is never observed in *Arabidopsis* where PINs are always present at the membranes facing neighboring cells (e.g., *12, 24, 25*). This PIN polarization in *Physcomitrium* strongly reduces intracellular auxin concentration by pumping a portion of auxin out of the organ, permitting sustained cell divisions at a distance from the base (Fig. 5). When PIN function is abolished, auxin accumulates intracellularly (Fig. 2E). This decreases cell divisions, stimulates cell elongation and differentiation, resulting in narrower phyllids compared to wild-type (Fig. 5). Exogenous auxin applications further inhibit cell divisions and stimulate cell elongation (especially in the absence of PINA and PINB) indicating that higher auxin concentrations in phyllids have increasingly stronger effect on their development, as observed in moss cultures grown with varying auxin concentrations (*41*).

**Fig. 5.**
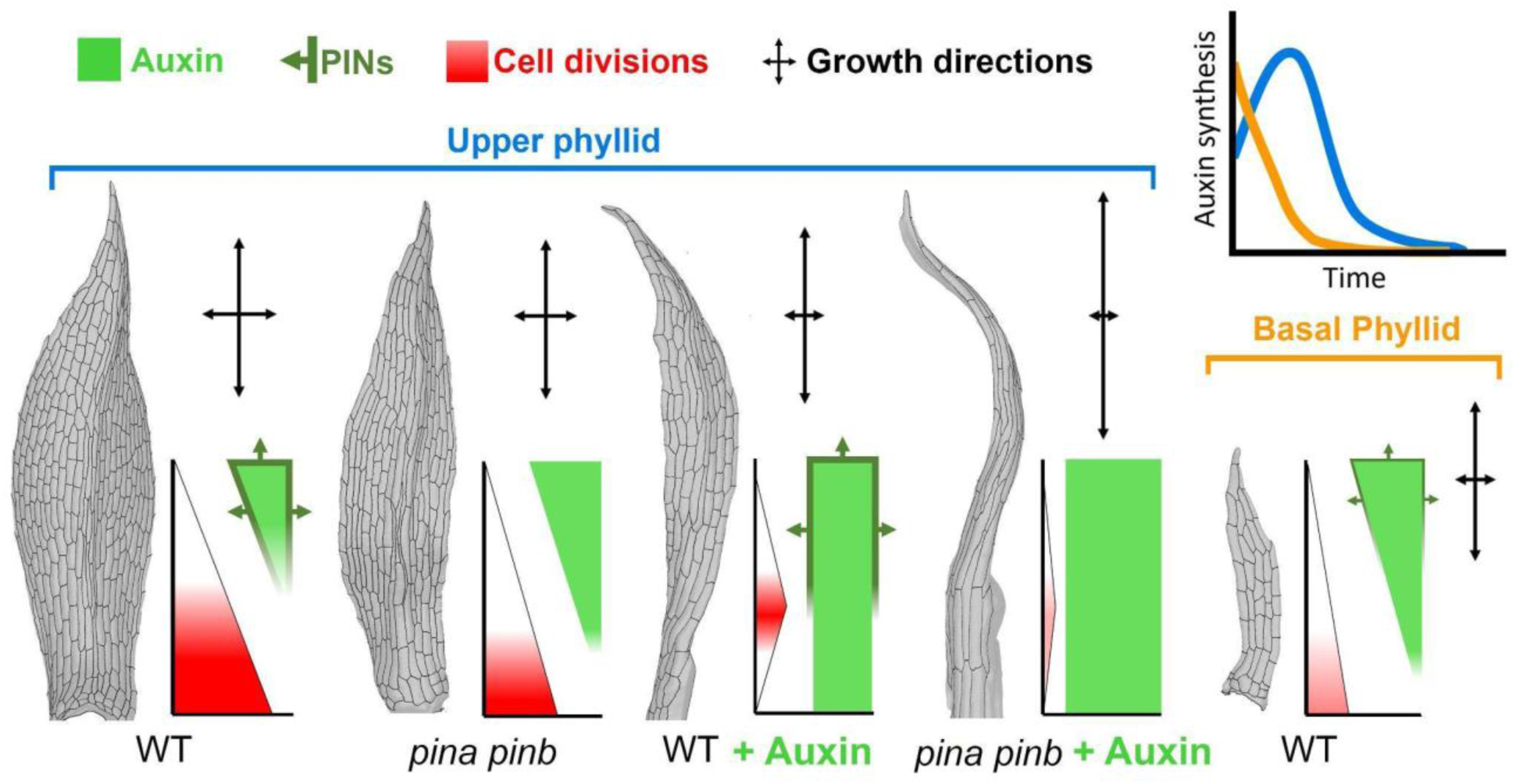
How auxin controls phyllid shape. By default, cell divisions are restricted to the organ base while auxin is produced at the tip and diffuses basipetally. Auxin inhibits cell divisions and promotes cell differentiation and elongation. In the wild-type upper phyllid, the activity of PIN auxin exporters maintains relatively low intracellular auxin levels, thereby allowing cell divisions to occur both relatively far from the organ base and over a longer period to produce broad phyllid. Eliminating PIN transporters raises intracellular auxin levels, enabling auxin to travel further away from the tip; consequently, cell divisions are reduced, and cell elongation is stimulated, yielding a narrower organ. External auxin application in the wild type overrides the basipetal cell division gradient. Cells at the base, which never express PINs, respond strongly to the exogenous auxin by undergoing early differentiation and elongation, while cells juxtaposed to the PIN-expressing region keep dividing for some time, ultimately yielding an even narrower phyllid. In the absence of PINs, auxin treatment quickly suppresses cell divisions, triggering strong elongation across the entire phyllid and yielding to the formation of rod-shaped organs. In the basal phyllid, auxin production peaks during initiation, causing an early transition from cell division to cell differentiation, and ultimately the development of a small organ with few cells.

Auxin levels in the midrib appear higher that in the phyllid blade (Fig. 2, D and E) which could suggest that auxin is involved in midrib formation and the midrib could transport auxin in *Physcomitrium*. In *Arabidopsis*, auxin is transported by polarized PIN efflux carriers (*46*), and PINs and auxin are upregulated in the developing pre-vascular tissue (*60*). While both PINA-GFP and PINB-GFP signals are detected in the moss midrib, in contrast to *Arabidopsis* PIN1, they are apolar and appear only 1.5-2 days after midrib development begins (Fig. 2b; fig. S4). Importantly, in mutants lacking a midrib such as *lateral suppressor 1* (*pplas1*) and *pplas2*, the phyllid and cell morphology remains similar to wild-type (*61, 62*). Together, these results suggest that the midrib plays only a minor role in auxin distribution throughout the phyllid and in determining final organ shape.

Exogenous auxin treatments further demonstrate the role of PINs in orchestrating gradients of cell growth and divisions. In wild-type phyllids, basal cells that lack PIN expression and are typically the last to differentiate, show a stronger response to auxin and undergo premature elongation and differentiation (Fig. 5). This effect is enhanced in *pin* mutants, reinforcing the idea that PINs prevent cell differentiation by keeping intracellular auxin levels relatively low. Time-lapse imaging of basal phyllids supports this model, showing that early *TARA* activity and limited PIN expression likely contribute to their smaller size by triggering precocious differentiation compared to adult phyllids (Fig. 5).

Unlike in angiosperms, we propose that *Physcomitrium* PINs may not be involved in long-range polar auxin transport during early phyllid development but rather in reducing intracellular auxin concentration by pumping auxin out of cells. This finding suggests that PIN function has diverged between angiosperms and mosses. Remarkably, the single PIN protein from the streptophyte alga *Klebsormidium flaccidum* is also distributed laterally and lacks specific polar localization when expressed in tobacco and moss cells (*63*). This discovery has led to the proposal that PIN’s ancestral role may have been to export auxin into the environment. Consistently, most of the auxin in moss cultures appears to accumulate in the medium (*64*). Thus, *Physcomitrium* PIN localization patterns could reflect an ancestral state in the green lineage. Moreover, this implies that intracellular auxin movement in phyllids may rely on alternative mechanisms, such as plasmodesmata-mediated diffusion. Symplasmic auxin movement has been implicated in *Arabidopsis* root development, hypocotyl tropism, and leaf venation (*53–56*), and auxin has been also shown to move via plasmodesmata in *Arabidopsis* leaf epidermis (*29*). In *Physcomitrium*, symplasmic auxin movement may be sufficient to regulate branch patterning in leafy shoots (*52, 53*) and numerous plasmodesmata are present in the developing phyllids in *Physcomitrium* (fig. S6). Further research is needed to determine whether this mechanism is sufficient alone to regulate developmental patterning in moss (*65*).

Another key difference between angiosperms and mosses is the degree to which auxin regulates organ development. In *Arabidopsis* leaves, exogenous auxin clearly accelerates cell growth and cell differentiation, but has limited effects on intrinsic developmental gradients (*12, 28*). In contrast, exogenous auxin treatments in *Physcomitrium* can override global basipetal gradients of cell division and growth, and in *pin* mutants, this leads to a dramatic perturbation of organ development, exemplified by the formation of rod-shaped phyllids. Overall, this underscores the role of auxin in influencing the final phyllid shapes.

Together, our findings in *Physcomitrium* and previous work in *Arabidopsis* and other angiosperms reveal several shared principles of planar organ morphogenesis. First, organ development is characterized by an initial phase of cell divisions followed by a transition to cell expansion and differentiation, coordinated by global positional cues (*12*, *15–19*). Second, auxin serves as a key positional cue regulating the transition from cell division to differentiation, with distinct contributions in juvenile and adult organs (*19*). We conclude that the convergent evolution of angiosperm leaves and moss phyllids was driven by the repeated deployment of these deeply shared developmental principles, with lineage- and species-specific variations.

## ACKNOWLEDGEMENTS

We thank Teva Vernoux and Dolf Weijers for insightful discussions, Bernd Reiss and Miltos Tsiantis for help with initial moss transformations, Ken Kosetsu for providing the EF1α promoter, Guillaume Cerrutti and Jonathan Legrand for help with image analysis, Stéphanie Hallet for technical support, Enrico Scarpella for critical reading of the manuscript, and Elvis Branchini and Daniel Grégoire for help with figure preparations.

## AUTHOR CONTRIBUTIONS

Conceptualization: WL, RSS, YC, and DK

Methodology: WL, MD, A-LR-L, and DK

Computational modelling: BL, LC, and RSS

Data acquisition: WL, LM, YC, AB-Z,

Visualization: WL, MD, BL, LM, DK, RSS

Funding acquisition: DK, A-LR-K and RSS

Project administration: DK

Supervision: A-LR-K, RSS, YC, and DK

Writing – original draft: WL, RSS, YC, and DK

Writing – reviewing and editing: A-LR-K, LM, MD

## DATA AND CODE AVAILABILITY

The data used to quantify growth parameters, models, and starting templates with a copy of the MDX software are available to download from the Open Science Framework repository (https://osf.io/2hwe6/)

## SUPPLEMENTARY MATHERIALS

Materials and Methods Figs. S1 to S15 References (67–79) Movies S1 to S15

