## Supplementary Methods and Figures for "Evolution of moss leaf-like organs through variations in deeply conserved developmental principles"

**The PDF file includes:**

Materials and Methods

Figs. S1 to S14

References (67-79)

**Other Supplementary Materials for this manuscript include the following:**

Movies S1 to S16

### MATERIAL AND METHODS

#### Plant materials and growth conditions

The following *Physcomitrum patens* transgenic lines were used: *PpEF1α::acyl-YFP* (this study), *pina pinb* double mutant (*34*), *pR2D2-1* (*445*) *PpPINApro::PpPINA-GFP* (*35*), *PpTARA::GFP-GUS* (*44*). *PpEF1α::acyl-YFP* was transformed into a *pina pinb* double mutant. The *pina* and *pinb* mutations were introduced into the *pR2D2-1* transgenic line to *achieve pR2D2-1-pinapinb* using the CRISPR-Cas9. *EF1α::acyl-YFP, pina pinb* and *PpPINApro::PpPINA-GFP* are in the Gransden background, all other reporter lines are in the Reute background. Plants were cultivated *in vitro* in sterile conditions on BCDAT medium (*66*) with 0.7% agar in a growth chamber under continuous light (80 µmol $m^{-2}s^{-1}$) at 22°C with 60-70% relative humidity. Exceptionally, *PpR2D2-1* and *PpR2D2-1-pinapinb* were cultivated under a 16h light and 8h dark cycle.

#### Construction of transgenes and plant transformation

Constructs for plant transformation were generated using standard cloning techniques.

To construct *PpEF1α::acyl-YFP*, the *acyl-YFP* sequence (*67*) was first moved from the Gateway destination vector *pUBQ10::acyl-YFP* (*68*) to the *pENTR207* donor vector (Thermo Fisher) using BP Gateway reaction. The *acyl-YFP* sequence in *pENTRE207* vector was then moved to pT1OG moss transformation vector containing *PpEF1α* promoter (provided by Ken Kosetsu, NIBB) and PTA1 homologous recombination sites using LR Gateway reaction to obtain *PpEF1α::acyl-YFP*. Final constructs were verified by sequencing and transformed into the wild-type using PEG-mediated homologous recombination method (*66*).

The *pina* and *pinb* mutations in *the R2D2-1* transgenic line (*45*) were generated using the Gateway based CRISPR-Cas9 vector system (*70*, *71*). Four protospacers (corresponding to sgRNA) targeting the 5'UTR (position 198), the exon 1 (position 1086), the exon 4 (position 106) and the 3'UTR (position 140) of *PpPINA* (Pp3c23_10200V3.1) and four additional protospacers targeting the 5'UTR (position 205), the exon 1 (position 1023), the exon 4 (position 122) and the 3'UTR (position 203) of *PpPINB* (Pp3c24_2970V3.1) were designed using the CRISPOR software (*72*). For each targeted gene, the four complementary protospacer oligos (Reverse and Forward pairs) were annealed and then inserted downstream of an U6 promoter, into the *BsaI* site of the following *pENTR* vectors: *pENTR-PpU6P-sgRNA-L5L4*, *pENTR-PpU6P-sgRNA-L1R5*, *pENTR-PpU6P-sgRNA-R4R3*, and *pENTR-PpU6P-sgRNA-L3L2*. Co-transformation of *PpR2D2-1* protoplasts with all the entry vectors that carry the *PpPINA* and *PpPINB sgRNAs* together with the *pZeo-Cas9-gate* destination vector that contains the Cas9 enzyme coding sequence were performed as described before (*67*). After antibiotic selection, the resistant clones were genotyped by PCR using primers external to the coding sequence of *PpPINA* and *PpPINB*. For *PpPINA*, the expected size for the endogenous locus size was of 3765 bp and we found large deletion (more than 1500 bp between Exon 1-1086 and 3'UTR-140 sgRNAs) for clones 25 and 29 using the primers P43 and P44. For *PpPINB*, the endogenous locus is of 3792 bp and we also found large deletion in clones 25 and 29 (more than 3000 bp between 5'UTR-205 and 3'UTR-203 sgRNAs) using the primers P47 and P48.

**Gene expression data**

*TAR* and *YUC* gene expression data were retrieved from the MAdLandExpression database (https://peatmoss.plantcode.cup.uni-freiburg.de/easy_gdb/index.php). These data were originally published in (*72-73*).

**Confocal microscopy**

Zeiss LSM800 or LSM980 upright confocal microscope with diode lasers and 25X or 40X immersion objectives (1 NA, Apochromat) were used for imaging. Confocal z-stacks were acquired with 0.5-1 μm distance in z-dimension at 1024 x 1024 for the young leaf primordium and 512 x 512 resolution for leaves at later developmental stages. YFP from the membrane marker and GFP from the reporter lines were excited with a 488 nm diode laser and the signal was collected at 400-614 nm and 500-550 nm respectively. Autofluorescence was simultaneously collected at 656-700 nm.

Airyscan mode was used to detect the detailed localization of PpPINApro::PpPINA-GFP in the young upper phyllids. Samples were flipped and images for both abaxial and adaxial sides were then merged together in MorphoGraphX software (*74*).

VENUS and TdTOMATO signals from *pR2D2-1* and *PpR2D2-1-pina pinb* reporter lines were collected as described before (*45*). Briefly, The *PpR2D2-1* and *PpR2D2-1 pina pinb* was imaged with a Zeiss LSM980 confocal microscope and a water immersion objective lens of 25X. The signals of DII (TdTomato) and mDII (VENUS) were imaged sequentially in order to avoid cross-talks.

For time-lapse imaging of the upper phyllids, gametophores with newly emerged 12th phyllid were selected from plates 18-days old. The neighboring phyllids were gently removed with fine tweezers to expose the target phyllid for imaging. The same phyllid was imaged every 0.5 days for about 6 days until maturity. If necessary, gametophores were tilted before the scanning to expose the abaxial side of the phyllid for imaging. Multiple overlapping stacks were taken to cover the samples exceeding the objective field of view. Plant samples were immersed in sterile water or chemical solutions during imaging. After each scanning, the plants were rinsed three times with sterile water, drained with tissue paper and returned to the original growth condition in the growth chamber. A minimum of three independent timelapse series were acquired for each experiment.

**Auxin treatment**

Auxin treatment was performed with 1 M 1-Naphthaleneacetic acid (NAA, Sigma) solution. Plants were submerging samples each 12 h in water with specified chemical solution during imaging. Treated samples were rinsed three times with sterile water after imaging and put back to the growth chamber.

**Electron microscopy**

For plasmodesmata visualization, the samples were processed following a previously described procedure (*75*). Dissected phyllids at around 2.5 – 3.5 days after initiation were fixed overnight at 4 °C in 2% glutaraldehyde and 2% formaldehyde (v/v) at pH 6.8, then post-fixed in 1% (v/v) osmium tetroxide for 2 h at room temperature. The samples were counterstained for 1 h with 2% uranyl acetate and embedded in low-viscosity Spurr resin. Ultrathin sections (70 nm) were cut along the longitudinal axis of the phyllids using a diamond knife on a Leica EM UC7 ultramicrotome (Leica-Reichert, Bensheim, Germany) and mounted on formvar-coated copper grids. Sections were subsequently stained with uranyl acetate and lead citrate and examined with a Hitachi HT7700 transmission electron microscope (Hitachi, Tokyo, Japan) operated at 80 kV.

**Image processing and analysis**

Confocal images were processed using MorphoGraphX software (*39*, *75*).

For volumetric analysis, 3D segmentation stacks were generated by applying the "gaussian blur" function twice before segmentation with the "ITK Watershed Auto Seeded" function with the threshold set to around 1500. The clipping plane was utilized to check through the samples to fix the wrongly segmented cells. Over-segmented labels were fused together manually. 3D meshes were then extracted from the volumetric stacks by "Marching Cubes 3D" function with 1 μm cube size followed by three smooth passes. "Grab label from other surface" and "Pick label in mesh1 and fill parent in mesh 2" tools were used to correlate cells with their parent labels. The clone identities were manually attributed to cells just derived from the leaf apical cell and propagated to later time points using cell lineage information. The boundary of the midrib is manually defined based on its multi-layer structure.

For 2.5D (curved surface) segmentations, stacks were "gaussian blur" twice before "edge detect" with a threshold between 6000 and 13000 followed by "edge detect angle" with a threshold between 4000 and 8000. 2.5 D meshes were then created using 5 um cube size, subdivided and smoothed 2-3 times before projecting signal (4-6 m away from the surface). The meshes were then manually segmented.

To project the signal intensity of *PpPINApro::PpPINA-GFP-1*, surfaces were first extracted from the autofluorescence images using procedures mentioned above. The signal from

*PpPINApro::PpPINA-GFP-1* at the range of 4 to 8 m was averaged and projected onto its corresponding meshes using the "Project Signal" function in MorphoGraphX software with parameter "Use absolute" set to "Yes" and "Max Signal" set to 40000.

Parent cells were manually identified and labeled between each two consecutive imaging. The cell lineage between multiple days was computed based on the parent labels. The clone identities were manually labeled as in volumetric analysis.

Cellular quantification is based on cell lineage information and the topology between neighboring cells. Area expansion calculated the relative area increase from the parent cell to its daughter cells in the next time point ([Total cell area of daughter cells at Ti+1/Area of the parent cell at Ti-1] 100 %). Cell division computed the number of divisions occurring in each parent cell which equals the number of daughter cells in the next time point minus one.

A small group of cells was manually labeled at the base of the leaf to define the leaf base region, ensuring that the selected boundary was perpendicular to the main leaf axis. This region was then used to compute distances from the base using the "Cell Distance" function in MorphoGraphX (with "Wall Weights" set to "Euclidean" to calculate absolute distance. The relative distance from the leaf base was achieved by normalizing the data with the maximum distance at the same time point.

The longitudinal (Kpar) and mediolateral (Kper) growth of phyllids was computed using the PDG analysis in MorphoGraphX and projected onto the custom directions (StretchCustomX = Kpar; StretchCustomY = Kper) extracted from the heat map of the absolute distance from the leaf base.

The new cell walls appearing in each time point were detected and highlighted by the "Select New Walls" function in MorphoGraphX. The orientation of the new wall (longitudinal or lateral) was manually identified and recorded.

To analyze the signal from *PpR2D2-1* and *PpR2D2-1-pina pinb*, confocal acquisitions were pre-processed using the ImageJ software (https://imagej.net/ij/) to select the region of interest (phyllid blades). The images were then processed using a computational pipeline described previously (*76*) which uses algorithms provided in the Python library timagetk (https://gitlab.inria.fr/mosaic/) to detect and quantify the signal from the nuclei.

**Model description**

**Model structure.** The model was implemented in the MorphoMechanX framework (*77–78*). The model was two-dimensional and consisted of structures at two distinct scales: a subdivision of the phyllid into cells, and a refinement of those cells into triangular finite elements; the refinement was constructed in such a way as to prevent elements from crossing cell boundaries. The cells were assigned growth parameters based on their type and position in the phyllid; these parameters were then passed down to their triangular refinement and used to determine the resultant growth of the tissue. Some cells in the model phyllid are capable of division; when a cell divided its finite elements were discarded, then the daughter cells were each triangulated into new elements. As residual stresses were released during the growth phase, this replacement caused no change in force distribution in the tissue.

**Growth and mechanics.** The finite elements are used in a growing finite element method (FEM) implementation that follows (*17*) but using 2D triangular 3-node membrane elements instead of

3D 6-node wedge elements. A linear St. Venant isotropic material model was used, with a Young's modulus of 100 MPa and Poisson's ratio of 0.3. Growth was implemented by increasing the size of the reference configuration of the elements at each growth step, with independent control of growth parallel (Kpar) or perpendicular (Kper) to a polarity direction. The rates of growth and the growth directions were specified at the cellular level, with polarity determined from the gradient of a distance field calculated from the apical cell. After each growth step, the mechanical equilibrium was found, vertex positions updated, and any residual stresses were released.

**Initial template.** A representative mesh at 1 day was projected flat into 2D, and the cell outlines were then extracted as polygons. These polygons were used as the model template. In the model, cells are again triangulated for the FEM, and this triangulation is updated when cells divide, together with a regularization of points along the edges to maintain reasonable triangle aspect ratios for simulation. The apical cell was given a specific identity. The bottom most vertices of the template were assigned Dirichlet conditions to prevent them moving in the longitudinal direction of the phyllid.

**Zonation and Cell Division.** The phyllid was divided into 3 zones. The attachment zone at the bottom had very slow growth. Above this was a basal zone determined by distance (in cells) from the base. In this zone cells were divided when they reached a threshold area. Division used cell polarity and picked the shorter of the dividing walls parallel or perpendicular to cell polarity. The apical cell was an exception: it divided into an alternating left-right pattern at 60°. As cells grew and moved out of the basal zone, they entered the differentiation zone where they stopped dividing. The model is parameterized in such a way as to allow the growth rates to be specified as a gradient within the zones, and to allow the zone sizes to change to match the phases of phyllid development and/or the influence of auxin. After each growth step, cell zones and locations were updated, and new parameters assigned based on their positional information.

**Midvien.** At a certain time point in the simulation, a midrib was specified by choosing cells within a certain proximity of the base and the central longitudinal axis. These cells would then be assigned different growth parameters, in particular a dramatic reduction in growth in the medial-lateral direction. As the simulation proceeded, the midrib identity was passed on to daughter cells.

### Statistical analysis

Data were extracted and analyzed using the Python 3.9 matplotlib library (https://matplotlib.org/stable/). Plots were generated using Excel, Python 3.9, Matplotlib, and Seaborn. In box plots, the central line represents the median, red dots indicate the mean, boxes span the interquartile range (1st to 3rd), and whiskers represent the 5th to 95th percentiles. The violin plots display the kernel probability distribution of the 5th to 95th percentiles of the data. Statistical significance was assessed with a two-sided Mann-Whitney U-test. To analyze growth patterns relative to the base of the leaf, the leaf was divided into 10 equal-length bins along its longitudinal axis. Median growth values within each bin were computed and visualized using a smooth spline interpolation, implemented via the make_interp_spline function from the scipy.interpolate module of the SciPy library. The variability in growth was represented by shaded regions around the spline curve, corresponding to the interquartile range (1st and 3rd quartiles) along the x-axis.

### SUPPLEMENTARY FIGURES

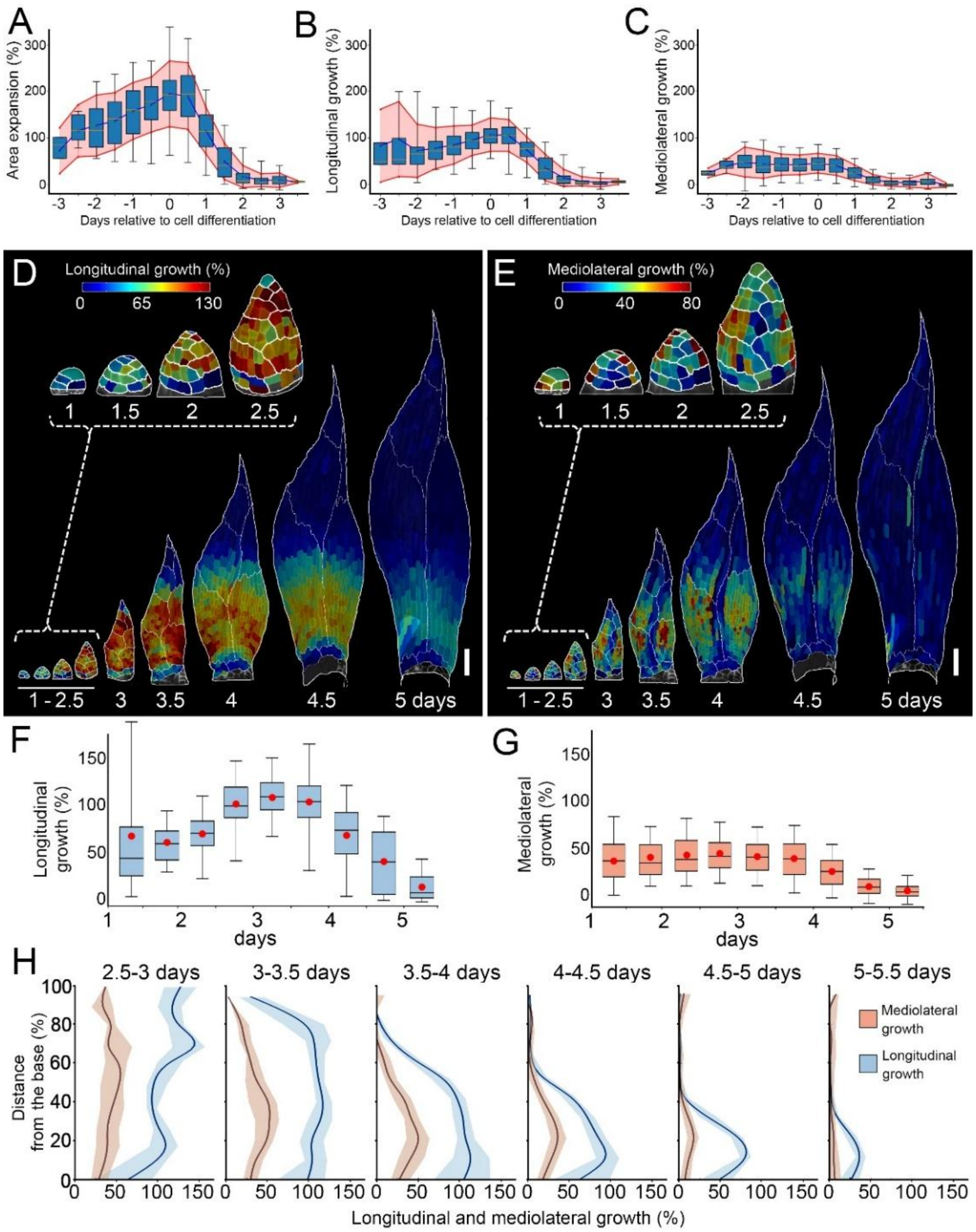


**Fig. S1. Mediolateral and longitudinal growth in the phyllid are independent from cell-lineage. (A-C)** Quantification of cell area expansion (A), cellular growth along longitudinal (B), and mediolateral (C) axis of the upper phyllid of *Physcomitrium patens* in days relative to cell differentiation (last division). **(D-E)** Heat-maps of cellular growth along longitudinal (D) and mediolateral (E) organ axis. Heat values are displayed at the earlier time point. **(F-G)** Quantifications of cellular growth along longitudinal (F), mediolateral (G) in the upper phyllid. Boxes contain the second and third quartile and whiskers 90 % of data. Lines represent the median and the red dots the mean (n=18, 46, 91, 226, 490, 799, 931, 939 and 938 cells at consecutive time points; three independent time-lapse series). **(H)** Quantification of cellular growth along longitudinal and mediolateral organ axis as a function of the normalized distance from its base. Shades contain the second and third quartile and lines indicate the median. Scale bars = 100 µm. Related to Fig. 1.

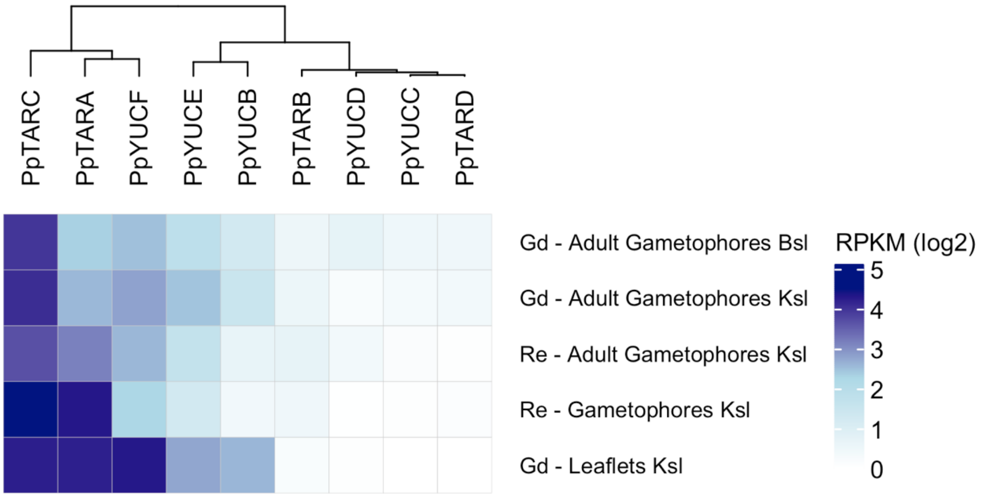


**Fig. S2. Expression of genes involved in auxin synthesis.** Heatmap showing expression levels of *PpTAR* and *PpYUC* genes in gametophores and phyllids from Gransden (Gd) and Reute (Re) ecotypes grown on KNOP solid medium (Ksl) or BCD solid medium (Bsl). All samples were normalized to RPKM.

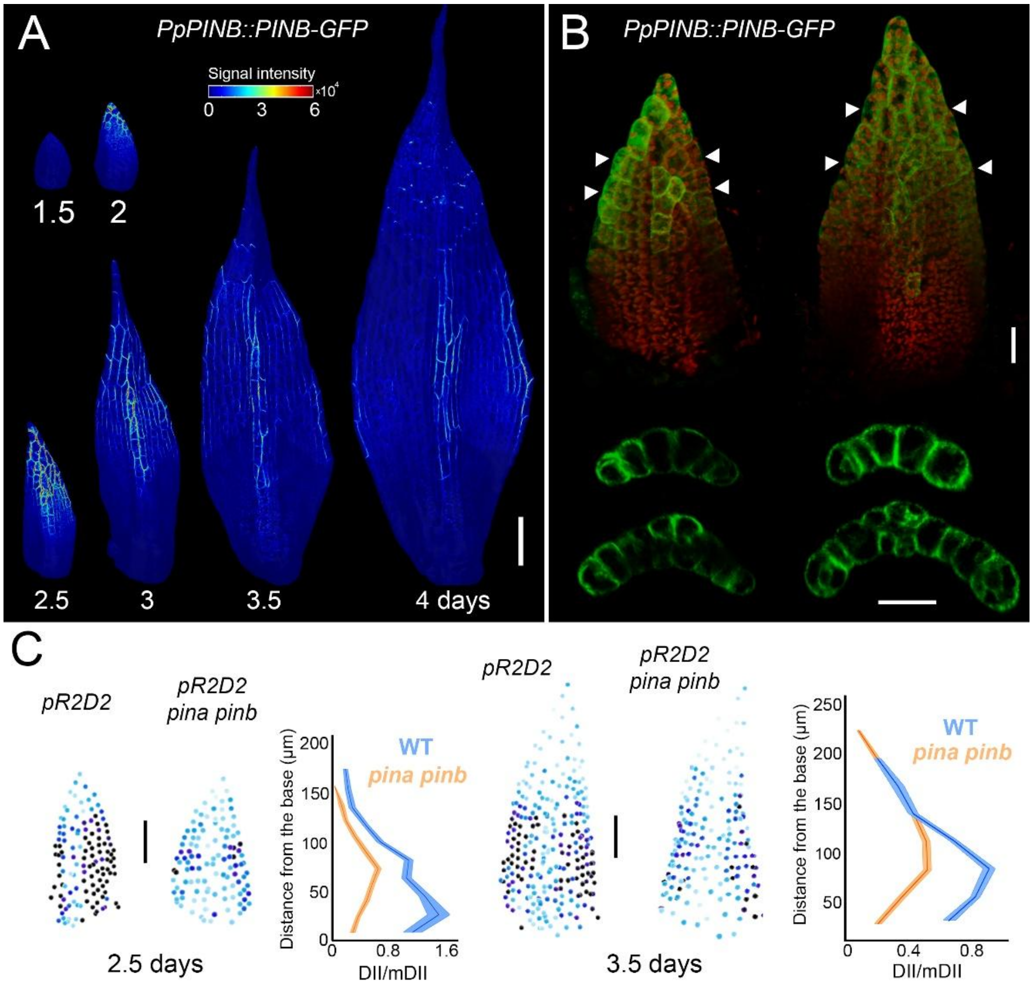


**Fig. S3. The localization of PINB auxin efflux carrier and the quantification of R2D2 signal distribution. (A)** Expression of *PpPINB::PINB-GFP* in the consecutive stages of upper phyllid development. Heatmap represents the intensity of PINB-GFP signal. **(B)** Maximal projections (top) and cross-sections (bottom) of confocal stacks with PINB-GFP signal in green and chloroplast autofluorescence in red. Arrowheads indicate the positions of the cross-sections**. (C)** Images of segmented nuclei in phyllids at 2.5 and 3.5 days. Each nucleus is color-coded according to the relative level of auxin sensing (DII/mDII signal ratio) (left) and quantification of DII/mDII signal ratio as a function of the absolute distance from the phyllid base. Shades contain the second and third quartile and lines indicate the median. Scale bar, 100 µm (A) and 20 µm (B). Related to Fig. 2.

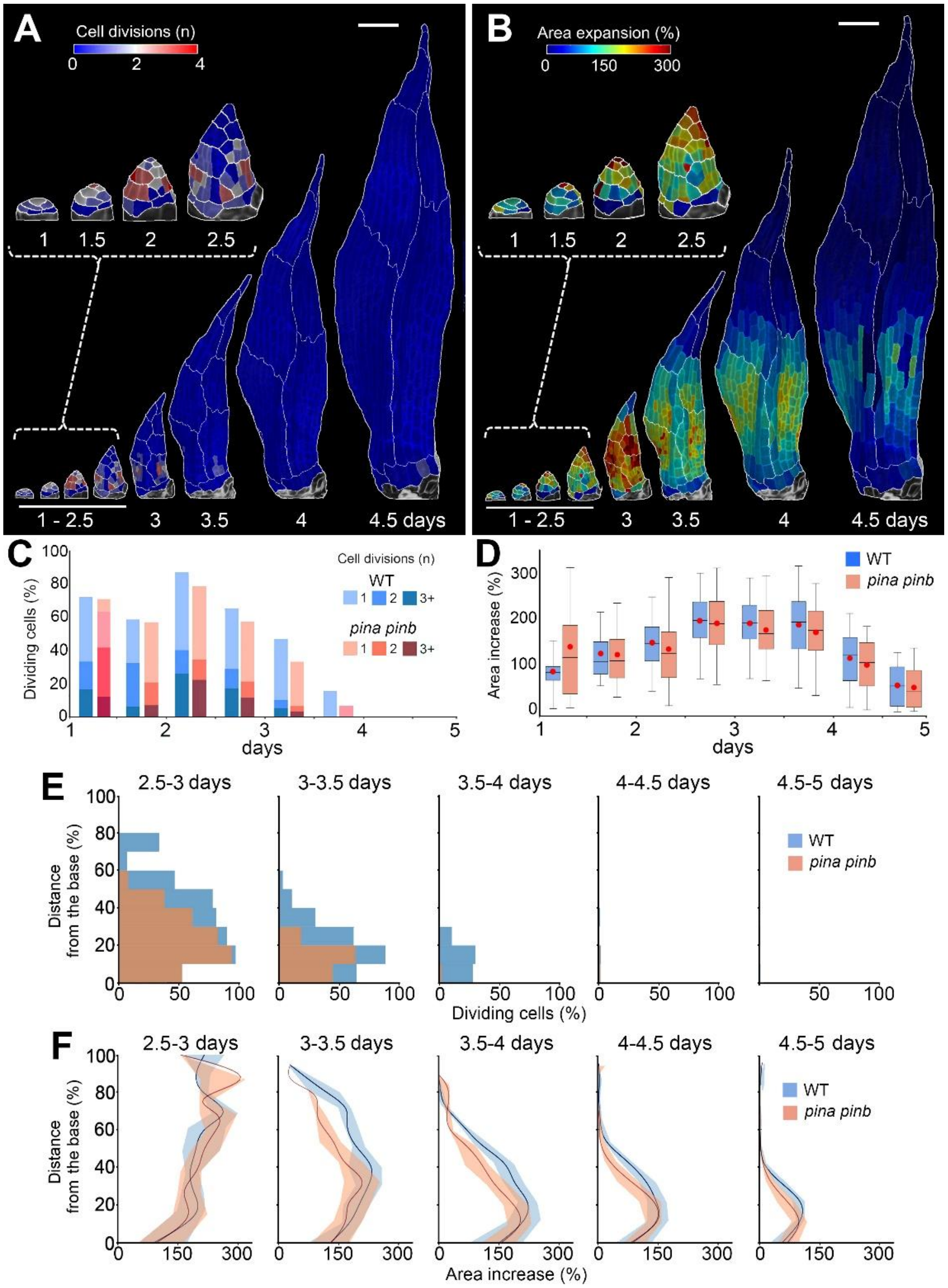


**Fig. S4. Removing PINA and PINB activity reduces cell division of phyllids. (A-B)** Heat-maps of cell divisions (A) and area increase (B) for the upper phyllid of *pina pinb* mutant. Heat values are displayed on the earlier time point. **(C-D)** Quantifications of cell divisions (C), and area increase (D) in the upper phyllid of the *pina pinb* mutant. Boxes contain the second and third quartile and whiskers 90 % of data (n=17, 37, 82, 195, 362, 510, 558 and 567 cells at consecutive time points; three time-lapse series). Lines represent the median and red dots the mean. **(E-F)** Quantification of the number of dividing cells (E) and area increases (F) as a function of the normalized distance from the organ base. Shades contain the second and third quartile and lines indicate the median. Scale bars = 100 µm. Related to Fig. 2.

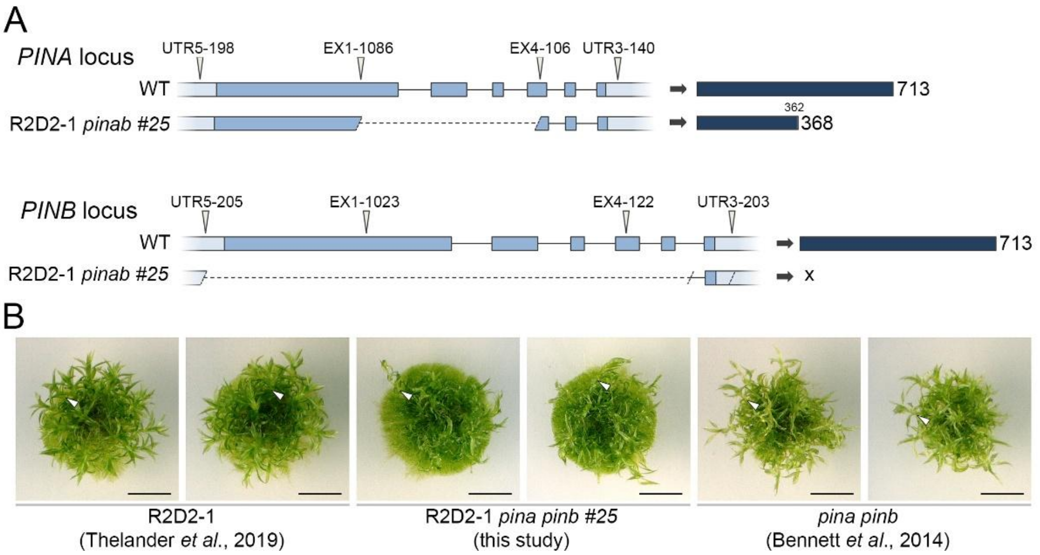


**Fig. S5. Genetic characterization and phenotyping of the R2D2-1 pinab line. (A)** Schematic representation of the WT *PINA* and *PINB* genomic locus showing the positions of four specific single guide RNAs (arrowheads) and the resulting genomic edits identified by sequencing in the R2D2-1 *pinab* #25 line (left). Blue boxes indicate exons; light blue boxes indicate 5′and 3′ untranslated regions (UTRs); introns are shown as solid black lines; dashed black lines indicate deleted regions of the genes. Predicted protein products resulting from an *in silico* translation of the edited genomic sequences (right). WT PINA and PINB proteins are 713 amino acids (AA) long. The mutant PINA protein is truncated to 368 AA, with a frameshift starting at position 362. **(B)** Photographs of representative 4-week-old clones of the R2D2-1 *pina pinb* #25 line generated in this study, compared to the original R2D2-1 genetic background. Gametophores (indicated by white arrowheads) in the R2D2-1 *pina pinb* #25 line display developmental alterations similar to those reported for the *pina pinb* double mutant in Bennett et al. (2014), confirming the efficiency of gene disruption. Note that the alterations of protonemal development observed in the R2D2-1 *pina pinb* #25 line were also present in the original R2D2-1 line and are therefore independent of PIN gene knockout. Scale bars, 0.5 cm. Related to Fig. 2.

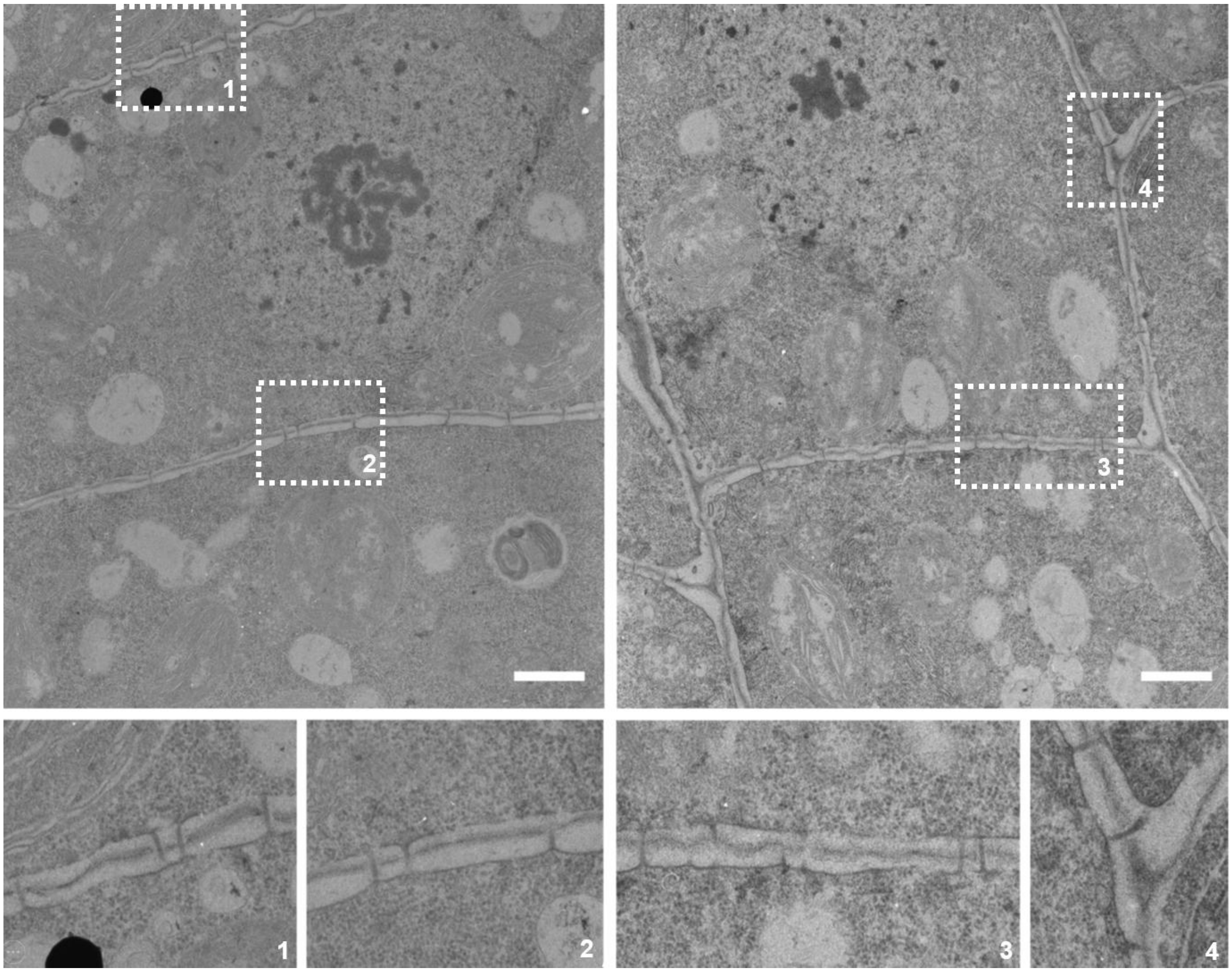


**Fig. S6. Visualization of the plasmodesmata in the anticlinal wall of the developing phyllids**. Representative transmission electron microscopy micrographs of phyllids at around 2.5-2.5 days after initiation. Insets indicate close up view of the plasmodesmata. Scale bars, 1 µm. Related to Fig. 2.

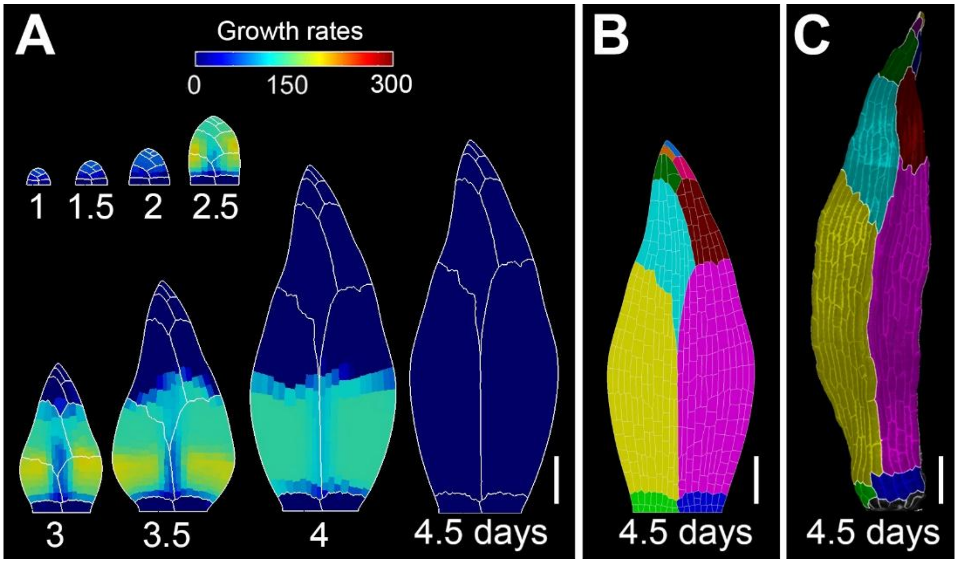


**Fig. S7. Model of phyllid growth with reduced cell division zone and the timing of cell proliferation. (A)** Model output colored by areal specified growth rate. **(B)** Resultant shape from the model colored by areal specified growth rate. **(C)** Fully developed upper phyllid of *pina pinb* mutant with colors marking sectors originating from the apical cell. Note that modelled phyllid is much shorter than double mutant. Scale bars, 100 µm. Related to Fig. 2.

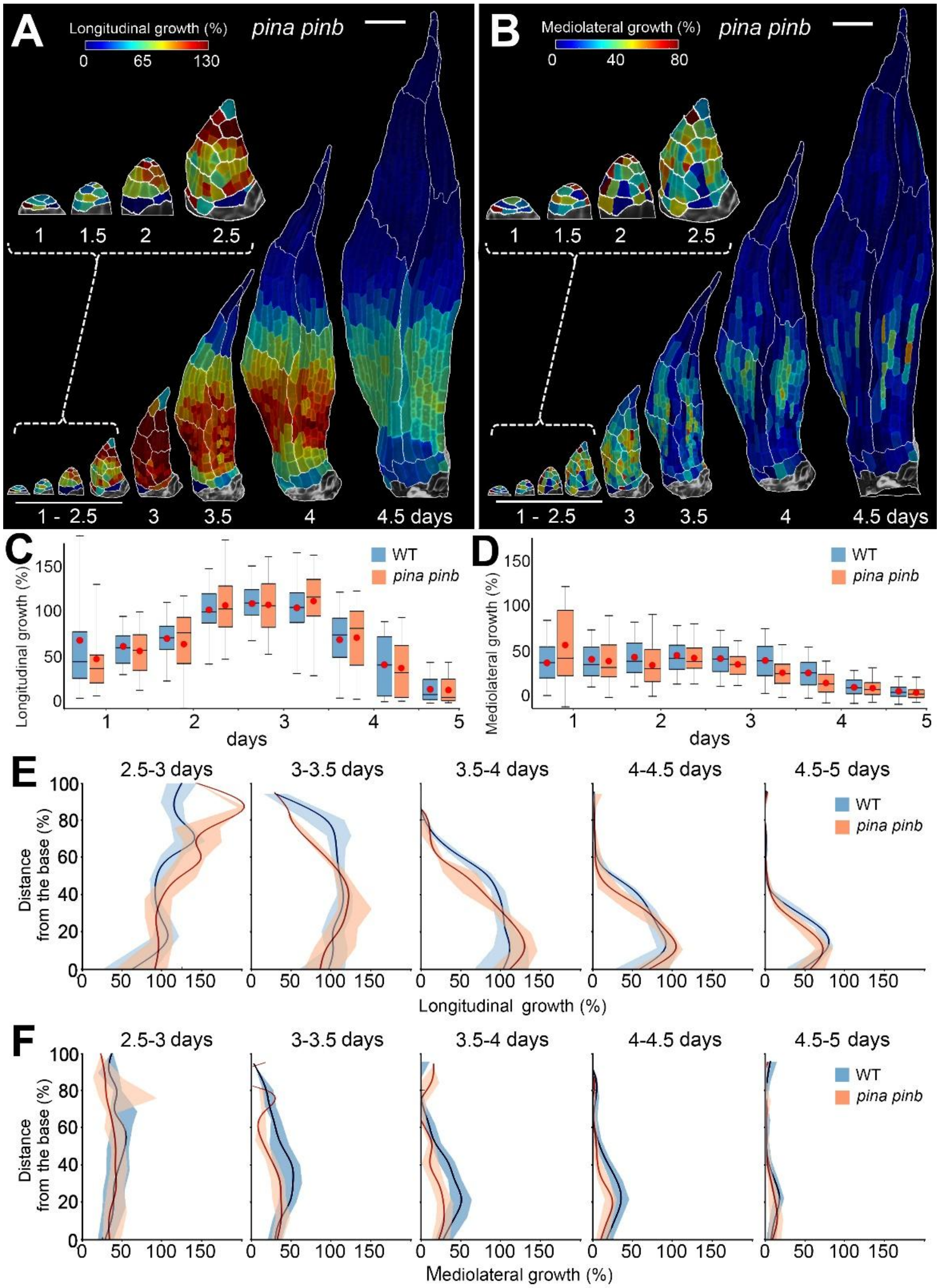


**Fig. S8. Removing PINA and PINB activity increases longitudinal growth and decreases mediolateral growth.** **(A-B)** Heat-maps of cellular growth along longitudinal (A) and mediolateral (B) axis of the upper phyllid of the *pina pinb* mutant. Heat values are displayed on the earlier time point. **(C-D)** Quantifications of cellular growth along longitudinal (C), medio-lateral (D) in the upper phyllid of the *pina pinb* mutant. Boxes contain the second and third quartile and whiskers 90 % of data (n=17, 37, 82, 195, 362, 510, 558 and 567 cells at consecutive time points; three time-lapse series). Lines represent the median and the red dots the mean. **(E-F)** Quantification of cellular growth along longitudinal (E), medio-lateral (F) as a function of the normalized distance from the organ base. Shades contain the second and third quartile and lines indicate the median. Scale bars = 100 µm. Related to Fig. 2.

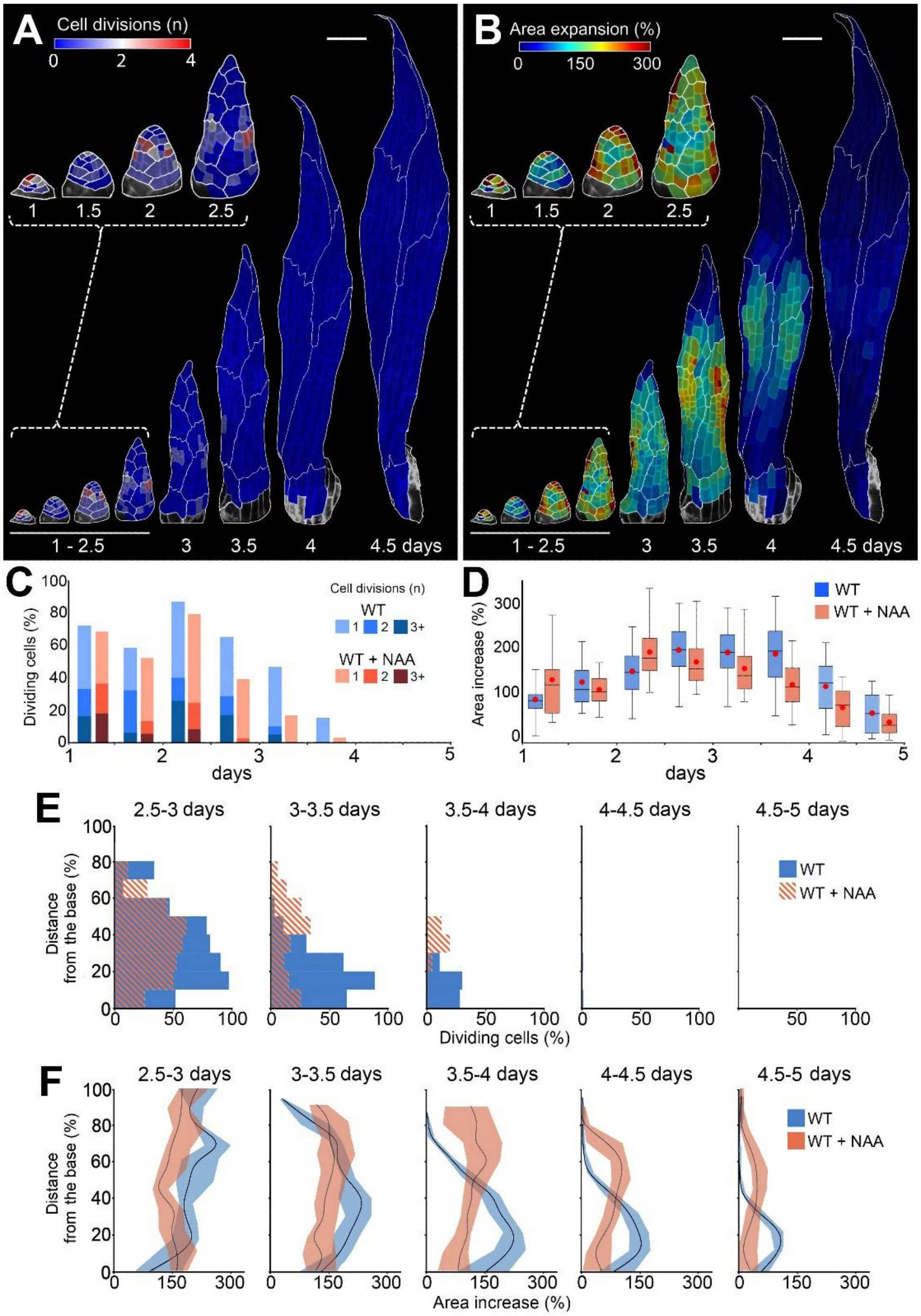


**Fig. S9. Auxin treatment perturbs basipetal gradients of cell proliferation and growth in the upper phyllid. (A-B)** Heat-maps of cell divisions (A) and area increase (B) for the upper phyllid upon NAA treatment. Heat values are displayed on the earlier time point. **(C-D)** Quantifications of cell divisions (C), and area increases (D) in the upper phyllid upon NAA treatment. Boxes contain the second and third quartile and whiskers 90 % of data (n=22, 52, 93, 193, 271, 321, 325 and 320 cells at consecutive time points; three time-lapse series). Lines represent the median and red dots the mean. **(E-F)** Quantification of the number of dividing cells (E) and area increases (F) as a function of the distance from the organ base. Shades contain the second and third quartile and lines indicate the median. Scale bars = 100 µm. Related to Fig. 3.

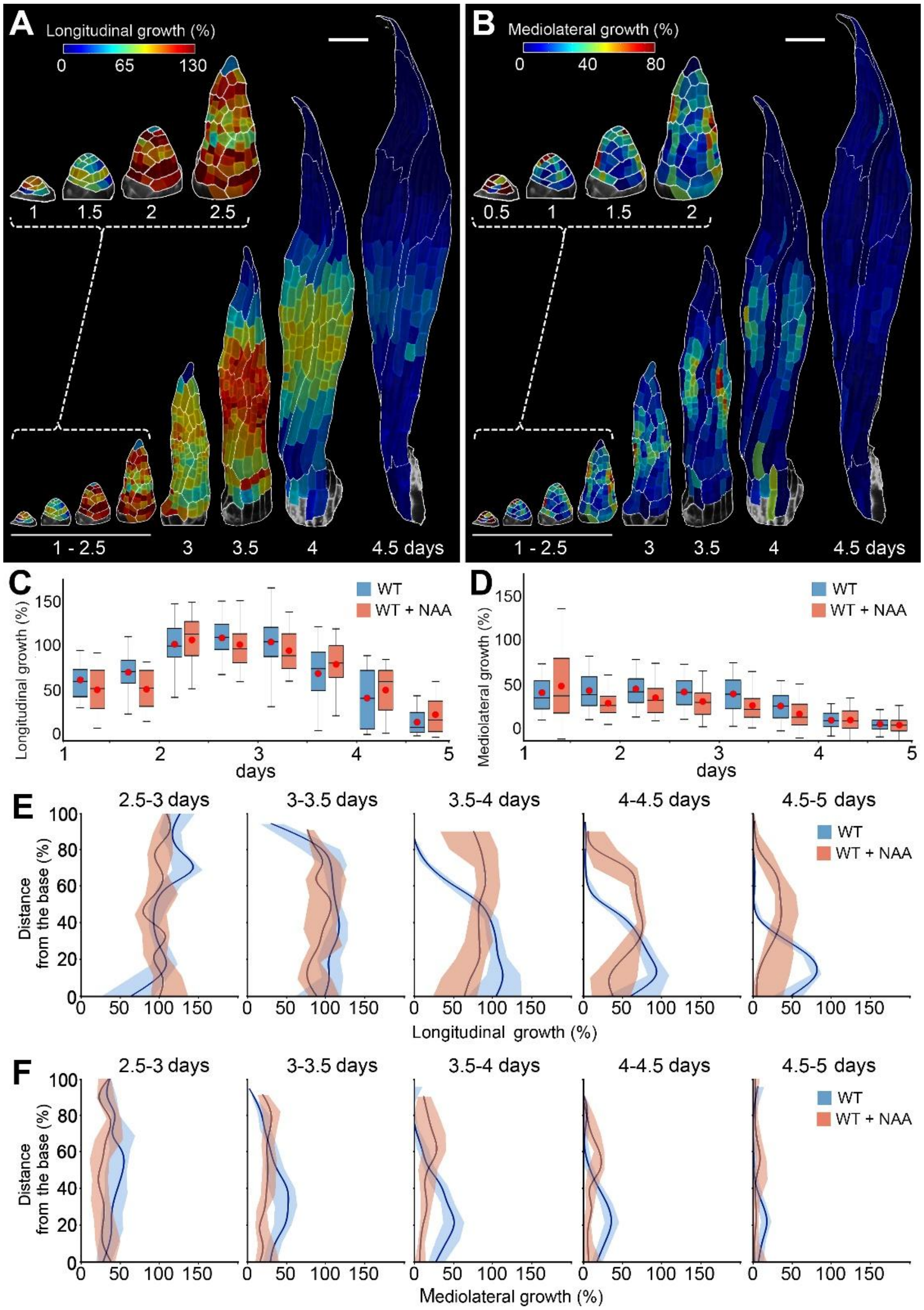


**Fig. S10. Auxin treatment perturbs basipetal gradients of growth in the upper phyllid. (A-B)** Heat-maps of cellular growth along longitudinal (A) and mediolateral (B) axis of the upper phyllid upon auxin treatment. Heat values are displayed on the earlier time point. **(C-D)** Quantifications of cellular growth along longitudinal (C), medio-lateral (D) in the upper phyllid upon auxin treatment. Boxes contain the second and third quartile and whiskers 90 % of data (n=22, 52, 93, 193, 271, 321, 325 and 320 cells at consecutive time points; three time-lapse series). Lines represent the median and the red dots the mean. **(E-F)** Quantification of cellular growth along longitudinal (E), medio-lateral (F) as a function of the normalized distance from the organ base. Shades contain the second and third quartile and lines indicate the median. Scale bars = 100 µm. Related to Fig. 3.

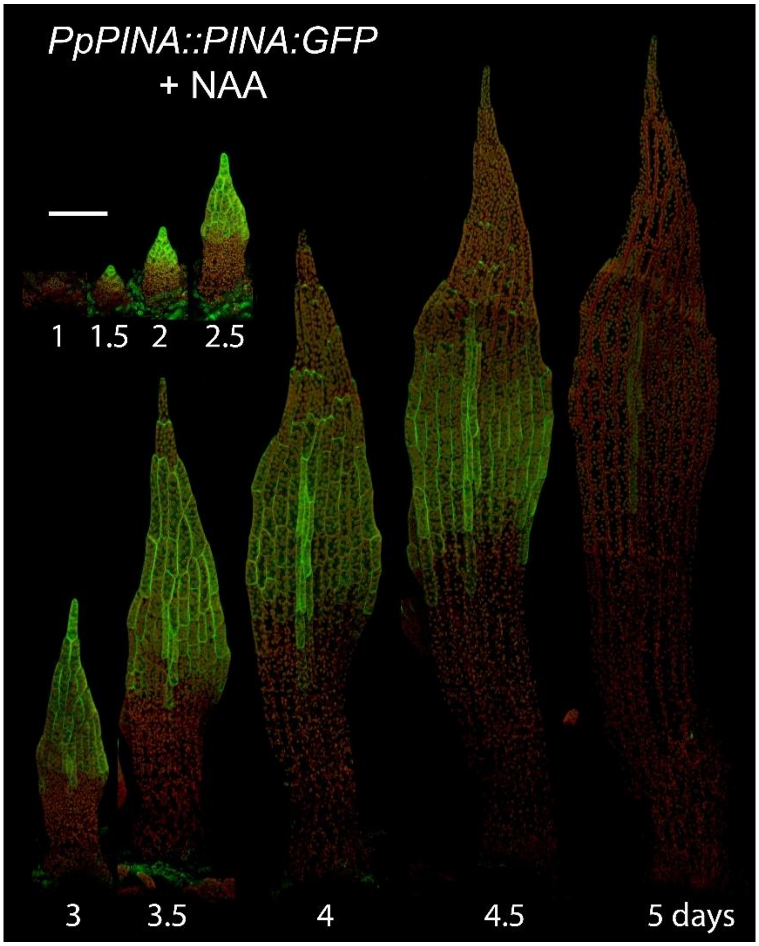


**Fig. S11. Auxin treatment does not affect the localization of PINA auxin efflux carrier.** Time-lapse imaging of the localization patterns of *PpPINA::PINA-GFP* during growth of the upper phyllis in the wild-type treated with NAA. PINA-GFP in green, autofluorescence in red. Scale bar, 100 µm. Related to Fig. 3.

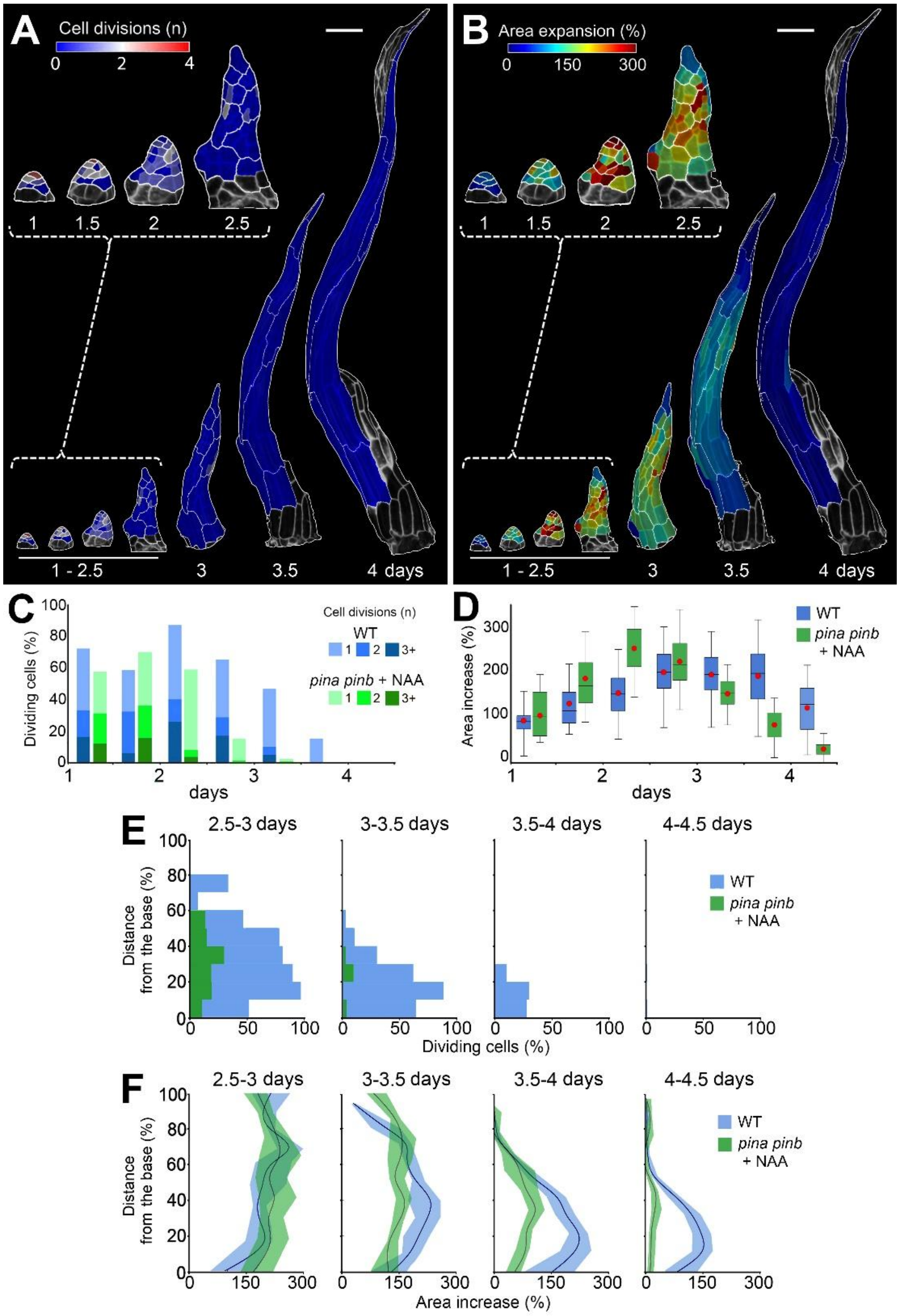


**Fig. S12. Auxin treatment of the *pina pinb* phyllid causes early cessation of cell divisions and growth. (A-B)** Heat-maps of cell divisions (A) and area increase (B) for the upper phyllid of *pina pinb* upon NAA treatment. Heat values are displayed on the earlier time point. **(C-D)** Quantifications of cell divisions (C), and area increases (D) in the upper phyllid in *pina pinb* mutant upon NAA treatment. Boxes contain the second and third quartile and whiskers 90 % of data (n=25, 52, 119, 201, 234, 239 and 237 cells at consecutive time points; three time-lapse series). Lines represent the median and red dots the mean **(E-F)** Quantification of the number of dividing cells (E) and area increases (F) as a function of the normalized distance from the organ base. Shades contain the second and third quartile and lines indicate the median. Scale bars = 100 µm. Related to Fig. 3.

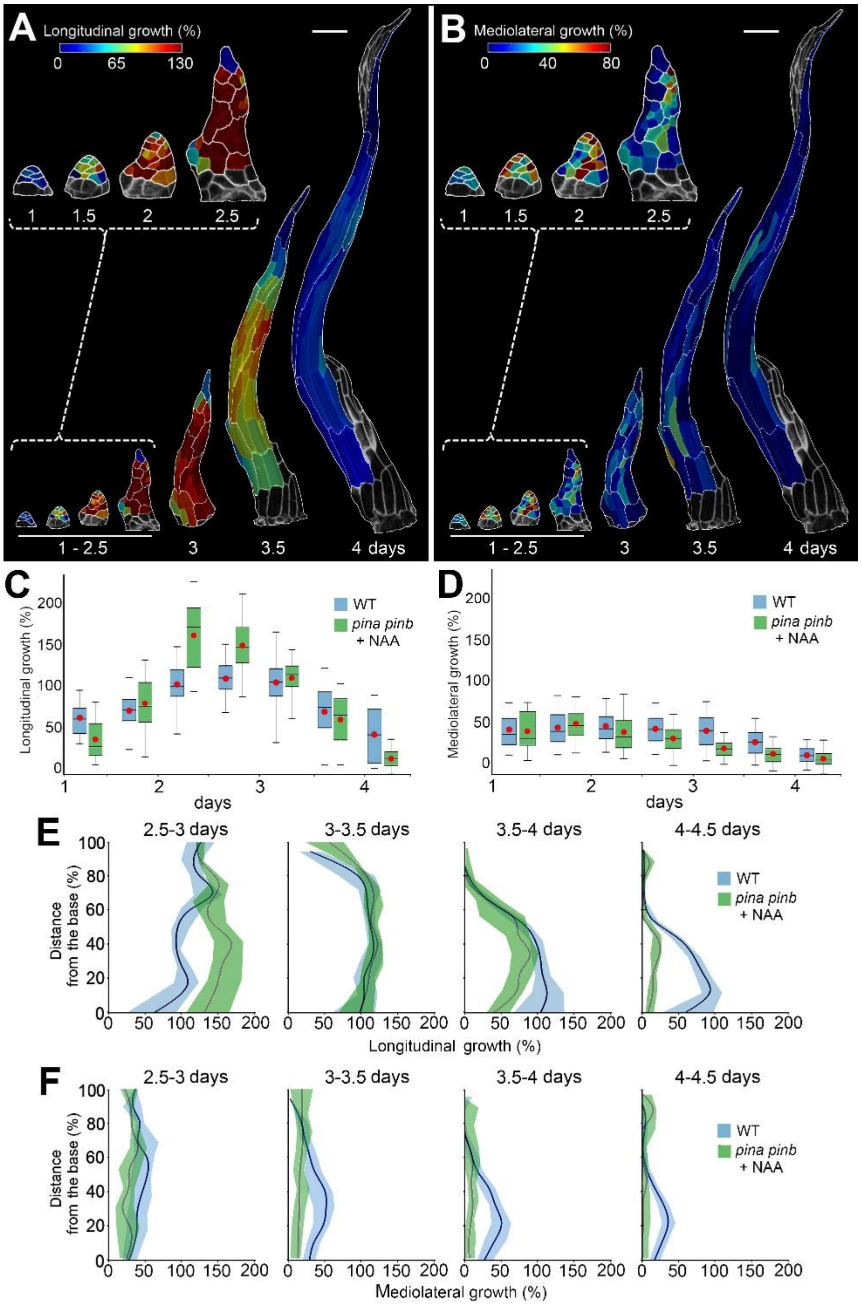


**Fig. S13. Auxin treatment of the *pina pinb* phyllid strongly increases longitudinal growth while blocking growth along the mediolateral axis. (A-B)** Heat-maps of cellular growth along longitudinal (A) and mediolateral (B) axis in the upper phyllid of the *pina pinb* mutant upon auxin treatment. Heat values are displayed at the earlier time point. **(C-D)** Quantifications of cellular growth along longitudinal (C), medio-lateral (D) in the upper phyllid of the *pina pinb* mutant upon auxin treatment. Boxes contain the second and third quartile and whiskers 90 % of data (n=25, 52, 119, 201, 234, 239 and 237 cells at consecutive time points; three time-lapse series). Lines represent the median and the red dots the mean. **(E-F)** Quantification of cellular growth along longitudinal (E), medio-lateral (F) as a function of the normalized distance from the organ base. Shades contain the second and third quartile and lines indicate the median. Scale bars = 100 µm. Related to Fig. 3.

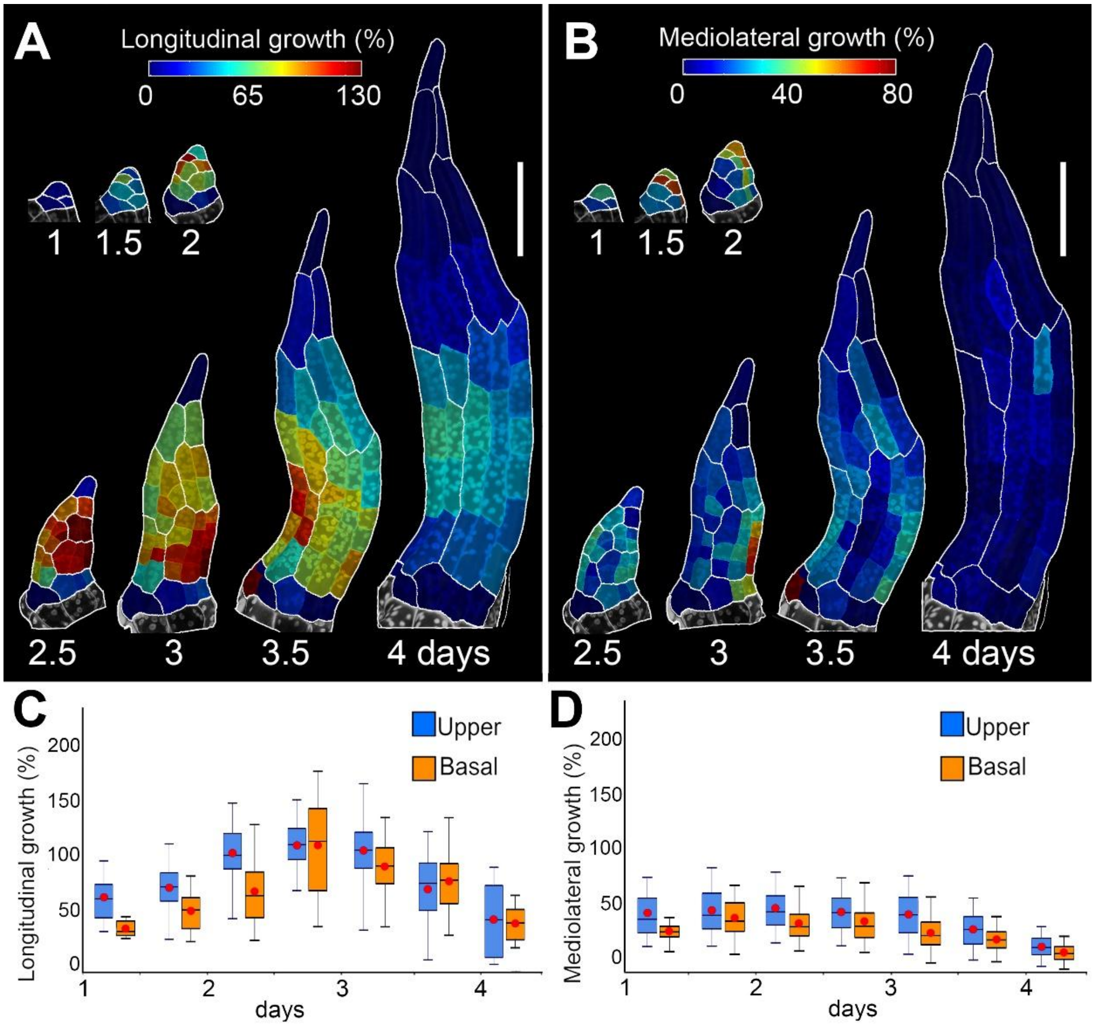


**Fig. S14. Growth of the basal phyllid is mainly longitudinal. (A-B)** Heat-maps of cellular growth along longitudinal (A) and mediolateral (B) axis in the basal phyllid. Heat values are displayed at the earlier point. **(C-D)** Quantifications of cellular growth along longitudinal (C), medio-lateral (D) in the basal phyllid. Boxes contain the second and third quartile and whiskers 90 % of data (n=15, 43, 94, 158, 238, 201 and 205 cells at consecutive time points; three time-lapse series). Lines represent the median and the red dots the mean. Scale bars = 100 µm. Related to Fig. 4.

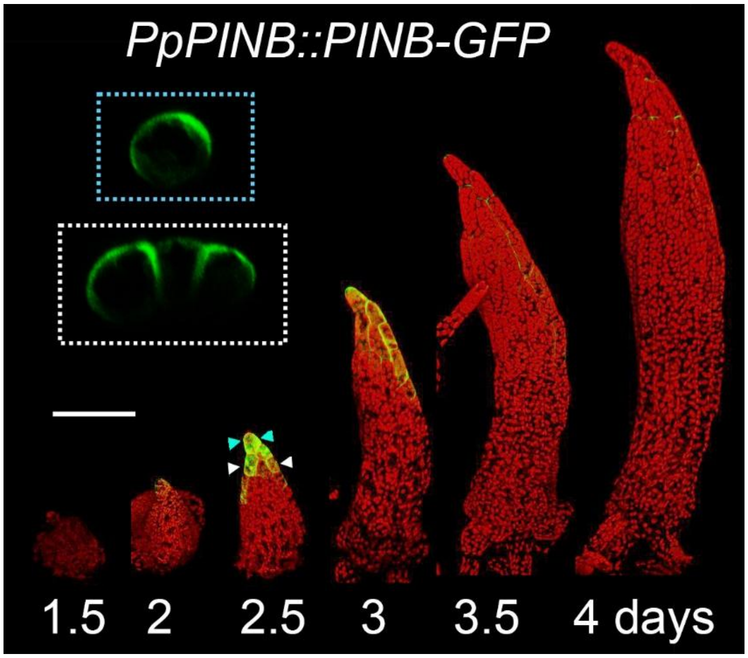


**Fig. S15**. The localization of PINB auxin efflux carrier. Expression of *PpPINB::PINB-GFP* in the consecutive stages of basal phyllid development. Maximal projections of confocal stacks with PINB-GFP signal in green and chloroplast autofluorescence in red. Insets: close-up view of the phyllid cross-sections. Arrowheads indicate the position of the cross sections shown. Scale bar, 100 μm. Related to Fig. 4.

### MOVIE LEGENDS

**Movie S1.** Cell division and area expansion in the wild-type upper phyllid of *Physcomitrium patens*.

**Movie S2.** Longitudinal growth and mediolateral growth in the wild-type upper phyllid of *Physcomitrium patens*.

**Movie S3.** Model of wild-type upper phyllid of *Physcomitrium patens*.

**Movie S4**: Cell division and area expansion in the *pina pinb* upper phyllid.

**Movie S5**: Model of *pina pinb* upper phyllid with reduced cell divisions compared to WT.

**Movie S6**: Longitudinal growth and mediolateral growth in *pina pinb* upper phyllid.

**Movie S7**: Model of *pina pinb* upper phyllid with reduced cell divisions and increased growth anisotropy as compared to WT.

**Movie S8**: Model of wild-type upper phyllid with cell divisions eliminated after Phase #1 and an increased growth anisotropy.

**Movie S9:** Cell division and area expansion in the wild-type upper phyllid treated with NAA.

**Movie S10**: Longitudinal growth and mediolateral growth in the wild-type upper phyllid treated with NAA.

**Movie S11**: Cell division and area expansion in the *pina pinb* upper phyllid treated with NAA.

**Movie S12**: Longitudinal growth and mediolateral growth in *pina pinb* upper phyllid treated with NAA.

**Movie S13**: Cell division and area expansion in the wild-type basal phyllid

**Movie S14**: Longitudinal growth and mediolateral growth in the wild-type basal phyllid.

**Movie S15**: Model of wild-type basal phyllid.
